# Tumor Neoantigenicity Assessment with CSiN Score Incorporates Clonality and Immunogenicity to Predict Immunotherapy Outcomes

**DOI:** 10.1101/2020.01.23.917625

**Authors:** Tianshi Lu, Shidan Wang, Lin Xu, Qinbo Zhou, Nirmish Singla, Jianjun Gao, Subrata Manna, Laurentiu Pop, Zhiqun Xie, Mingyi Chen, Jason J. Luke, James Brugarolas, Raquibul Hannan, Tao Wang

## Abstract

Lack of responsiveness to checkpoint inhibitors is a central problem in the modern era of cancer immunotherapy. Tumor neoantigens are critical mediators of host immune response and immunotherapy treatment efficacy. Current studies of neoantigens almost entirely focus on total neoantigen load, which simplistically treats all neoantigens equally. Besides, neoantigen loads have been linked with treatment response and prognosis only in some studies, but not others. We developed a Cauchy-Schwarz index of Neoantigens (CSiN) score to characterize the degree of concentration of immunogenic neoantigens in truncal mutations. Unlike simple neoantigen loads, CSiN incorporates the effect of both clonality and MHC-binding affinity of neoantigens when characterizing patient neoantigen profiles. By exploiting the clinical responses in 501 treated patients (mostly by checkpoint inhibitors) and the overall survival of 1,978 baseline patients, we showed that CSiN scores predict treatment response to checkpoint inhibitors and prognosis in melanoma, lung cancer, and kidney cancer patients. CSiN substantially outperforms prior genetics-based prediction methods of responsiveness. Overall, our work fulfilled an important gap in current research involving neoantigens.

**One Sentence Summary:** The quality of tumor neoantigens predicts response to immunotherapy

## Introduction

Most immunotherapies, including checkpoint inhibitors, benefit only a small subset of patients. For example, anti-PD-1 and anti-PD-L1 agents, which have demonstrated marked clinical benefit in various diseases, have overall response rates ranging from 10%-50% in melanoma and non-small lung cancer (*1*–*5*) and higher response rate in some other select tumor types, such as classic Hodgkin lymphoma (*6, 7*) (65-80%). Similarly, the efficacy of anti-CTLA4 agents ranges from 10-15% for objective response rates and <3% for complete response (*8*). Unfortunately, the field is still far from clearly understanding how to distinguish responders and non-responders to immunotherapy. All forms of immunotherapy, such as checkpoint inhibitors and neoantigen vaccines, seek to activate the host immune system to attack the tumor cells. These forms of immunotherapy have different modes of actions, but most are intended to mobilize the cytotoxicity of T cells in the patient. Neoantigens are the most potent targets of T cell responses (*9*–*13*) and the profiles of neoantigens in each patient are central to determining the responsiveness to immunotherapy treatment.

A major impediment of current research that seeks to correlate neoantigens with immunotherapy treatment response is that most studies have only considered whether a higher neoantigen/mutation load (namely, the total number of neoantigens or mutations) is correlated with better immunotherapy response. This simplistic approach misses the rich information contained in the whole repertoire of neoantigens and has been successful in only some studies (*2, 4, 14*–*18*), but not others (*19*–*24*). Neoantigens are associated with mutations that can be either truncal or subclonal. Some neoantigens are also more immunogenic than others. This rich information is not captured by the unsophisticated neoantigen/mutation load approach, but could be critical for understanding the responsiveness of cancer patients to immunotherapy treatment. For example, Miao *et al* showed that cancer patients with a high proportion of clonal mutations have better rates of checkpoint inhibitor treatment response (*25*). The only other study that we are now aware of and that has defined a more sophisticated neoantigen-based predictive metric is the neoantigen fitness model developed based on evolutionary modeling of patient neoantigen profiles (*16, 26*). This work considered the neoantigen-class I MHC binding affinity and only retained the top neoantigen from missense mutations with the highest binding affinity within each tumor clone. This metric demonstrated excellent predictive power for survival of patients after immunotherapy treatment in a few cohorts, however, its predictive values and prognostic values have not been widely evaluated.

In this study, we developed CSiN for a quantitative characterization of the delicate internal structure of patient tumor neoantigen profiles. We assembled 10 immunotherapy-treated patient cohorts and 6 patient cohorts with baseline survival information, both from public sources and also our own cohorts. In our unbiased analysis, CSiN achieved almost uniformly significant results on the checkpoint inhibitor cohorts and the baseline cohorts of immunogenic cancers, which is significantly better than previously reported neoantigen-based biomarkers. Taken together, our work filled an important “void” in immune-oncological research that resulted from overlooking neoantigen clonal structures.

## Results

### Constructing the Cauchy-Schwarz index of Neoantigen (CSiN) to characterize the fine structure of patient neoantigen repertoire

Some somatic mutations are truncal and other somatic mutations are subclonal. Truncal mutations are shared among more tumor clones (founding and derivative clones) and, if targeted by T cells through neoantigens, would likely cause a stronger cytotoxic effect. Subclonal mutations are unique to different clones, and if a subclonal mutation is in a clone with a larger clonal fraction, the neoantigens associated with this mutation are likely to have a stronger effect on the survival of the tumor cells than subclonal mutations associated with minor clones. Besides, each somatic mutation could generate a different number of neoantigens of different peptide lengths (8 to 11 amino acids for class I MHC-binding peptides, 15 amino acids for class II-binding peptides), with different registers (a sliding window of protein segments around the mutated position) and presented by different HLA alleles (class I and class II). Insertions and deletions usually generate a higher number of neoantigens per mutation than missense mutations as they lead to the translation of completely new segments of protein sequences (shown in **Supplementary Information**). We hypothesize that a favorable distribution of neoantigens and tumor mutations occurs when immunogenic neoantigens are concentrated on truncal mutations (**Fig. 1A**), and we developed CSiN to characterize this.

**Fig. 1.**
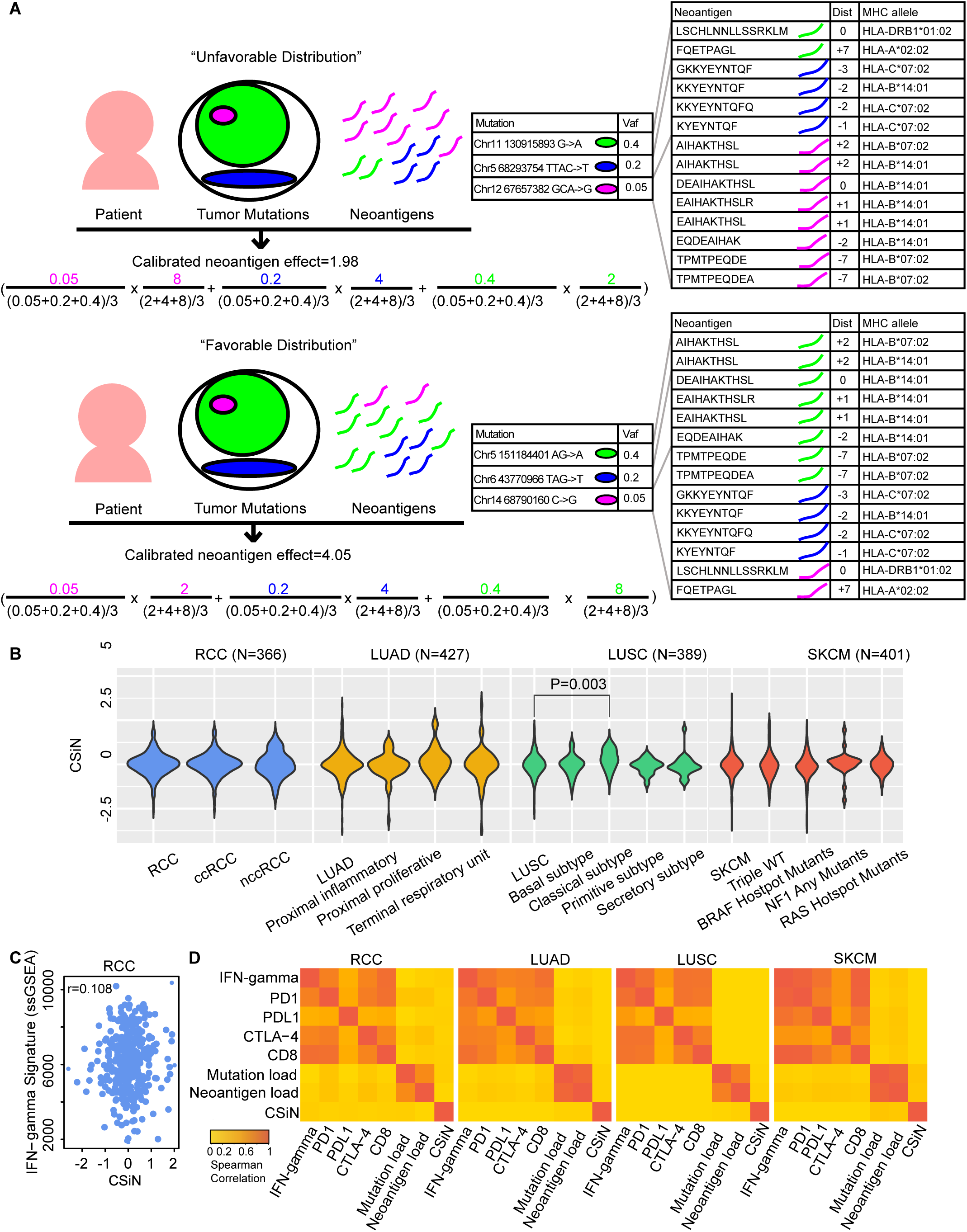
Motivation for CSiN. (A) Illustration showing the motivation of examining pairings of neoantigens and the tumor mutations with which they are associated. We demonstrated two fake patients, one with an unfavorable distribution and the other with a favorable distribution of neoantigens. But the mutations and neoantigens shown are taken from real data. The outermost circle indicates the whole tumor. Each circle indicates a population of tumor cells with certain mutations. Each different color indicates a distinct mutation, and the area of each circle indicates the proportion of cells having the mutation. For the formula, on the left of each multiplication sign “x”, is the normalized VAF, and on the right of each “x” is the normalized per-mutation neoantigen load. The colorings in the formula correspond to the tumor mutations shown above with the same colorings. The two bigger tables on the right show the neoantigen sequences, registers (“Dist”), and the HLA alleles for each neoantigen. For neoantigens of missense mutations, “Dist” refers to the distance between the altered amino acid and the left end of neoantigen; for neoantigens of insertions/deletions and stoploss mutations, “Dist” refers to the distance between the left end of the mutation and the left end of neoantigen. The “+” sign indicates the left end of neoantigen is on the right of the altered position and *vice versa*. (B) The distribution of the CSiN scores in the RCC, LUAD, LUSC, and SKCM cohorts. T-tests were used for comparison of CSiN scores between different subtypes of the same tumor cohort. (C) A scatterplot showing the relationship between CSiN and the expression level of the IFN-gamma signature in the RCC cohort. Spearman correlation between them is shown. (D) Heatmaps of the pairwise Spearman correlations of the CSiN, mutation load, neoantigen load, and the transcriptomics-based features are shown for the RCC, LUAD, LUSC and SKCM cohorts, which are calculated as in (D).

The core of CSiN is based on the mean of the product of the variant allele frequency (VAF) of each somatic mutation and the number of neoantigens from that mutation. When the mutations with higher VAFs are also the mutations that generate more neoantigens (favorable distribution), the product value will be larger (a higher CSiN score). Therefore, a higher CSiN conforms to a favorable neoantigen clonal structure. CSiN is so named because it bears analogy to the Cauchy-Schwarz inequality, which states the inner product of two vectors is maximal when they are in parallel (in the same ranked order). We also considered the immunogenicity quality of neoantigens. As there are currently no well accepted tools for predicting the potential of both class I and class II neoantigens to induce T cell responses (*27*), we used the binding affinity of peptides to MHCs, which has been shown to be one of the most important predictors of neoantigen immunogenicity (*28*). We set a series of cutoffs on the peptide-MHC binding affinity strengths predicted by the IEDB tools (*29, 30*) to generate several subsets of neoantigens with increasing stringency. We then calculated the means of product of VAF and per-mutation neoantigen load (neoantigens of both class I and class II, all HLA alleles, and all registers from one mutation) for each subset of neoantigens. The final CSiN score is the arithmetic mean of these sub-indices, in which neoantigens with better MHC binding affinity have higher weights (for details, please refer to **Supplementary Information**). The distributions of the CSiN scores in Renal Cell Carcinoma (RCC), lung adenocarcinoma (LUAD), lung squamous cell carcinoma (LUSC), and melanoma (SKCM), which are all immunogenic tumors (*31*), are shown in **Fig. 1B**.

We assessed whether CSiN characterizes unique information that is not captured by the commonly used mutation load and neoantigen load approaches (both classes, all possible lengths and all possible registers) or by candidate transcriptomic-based predictive biomarkers. For transcriptomic features, we examined the expression of *PD-1, PD-L1, CTLA-4 (32), CD8*, and an IFN-gamma gene signature (*33*). **Fig. 1C** shows a scatter plot of the CSiN scores with the activation (ssGSEA) of the IFN-gamma signature in the RCC cohort, which yielded a Spearman correlation of 0.067. **Fig. 1D** visualizes these correlations in heatmaps for pairwise comparisons of these biomarkers in each cohort, which shows that CSiN is an independent biomarker (Spearman correlation<0.1 for all comparisons). Further analyses, using the Pearson correlation, threshold comparisons, and mutual information, again demonstrated their independence (**Supplementary Information**).

### Better response to checkpoint inhibitors in immunogenic cancers is associated with higher CSiN scores

Next we investigated the implications of CSiN for checkpoint inhibitor treatment response. We analyzed the neoantigen profiles of melanoma patients on anti-CTLA-4 therapy from Van Allen *et al* (*34*). We observed patients with better response were more likely to have high CSiN (higher than median) than patients with worse response (P=0.009, **Fig. 2A**). We analyzed another cohort of melanoma patients on anti-CTLA-4 therapy from Snyder *et al* (*35*). We also observed that patients who received Durable Clinical Benefit (DCB defined as CR/PR/SD>6 month) had higher CSiN scores than patients with No Durable Benefit (NDB, SD<6 month/PD) (P=0.033, **Fig. 2B**). We analyzed a third cohort of melanoma patients (Riaz cohort) on anti-PD-1 therapy (*15*). We examined whether patients with better treatment response have higher CSiN scores. There is indeed a significant positive association in this cohort (P=0.037, **Fig. 2C**). We analyzed one more cohort (Hugo) of melanoma patients from Hugo *et al (14)*. **Fig. 2D** shows that there is an overall trend of patients with better response associated with higher CSiN scores (P=0.043). In clear cell RCC (ccRCC), we examined anti-PD-1/anti-PD-L1-treated ccRCC patients from Miao *et al (17)*. The same significantly positive association of higher CSiN scores with better response was observed (**Fig. 2E**) (P=0.036). We analyzed metastatic ccRCC patients treated with atezolizumab, an anti-PDL1 agent (Immotion150 cohort) (*36*). We found that there was a significant association of higher CSiN with better treatment response for T_eff_-high patients treated with atezolizumab (**Fig. 2F**, P=0.028 and **Supplementary Information**). In contrast, we did not observe this association for T_eff_-high patients treated with sunitinib (P=0.890). For patients with lower T_eff_ signature expression, there was no significant association for either atezolizumab or sunitinib. Non Small Cell Lung Cancer (NSCLC) patients (the Hellmann cohort) treated with PD-1 and CTLA-4 inhibitors were available from Hellmann *et al (37)*. Our analyses showed that PD-L1+ patients with Durable Clinical Benefit had higher CSiN scores than patients with No Durable Benefit (**Fig. 2G**, P=0.007), while this association is insignificant for patients of low PD-L1 expression. We examined another NSCLC cohort (the Acquired cohort) from Anagnostou *et al (2)* and Gettinger *et al (22)*. All of these patients achieved partial response, except for one patient who exhibited stable disease after initial checkpoint inhibitor treatment. All patients developed acquired resistance after 4 to 40 months. Interestingly, patients with sustained response were more likely to have higher CSiN scores than patients with short term progression (**Fig. 2H**) (P=0.015). Lastly, another cohort of NSCLC patients on anti-PD-1 therapies from Rizvi *et al (4)* was analyzed. In **Fig. 2I**, we showed that more DCB patients had higher CSiN scores than NDB patients (P=0.058). The False Discovery Rates for all the above mentioned cohorts fall under 10% (**Sup. Table 3**).

**Fig. 2.**
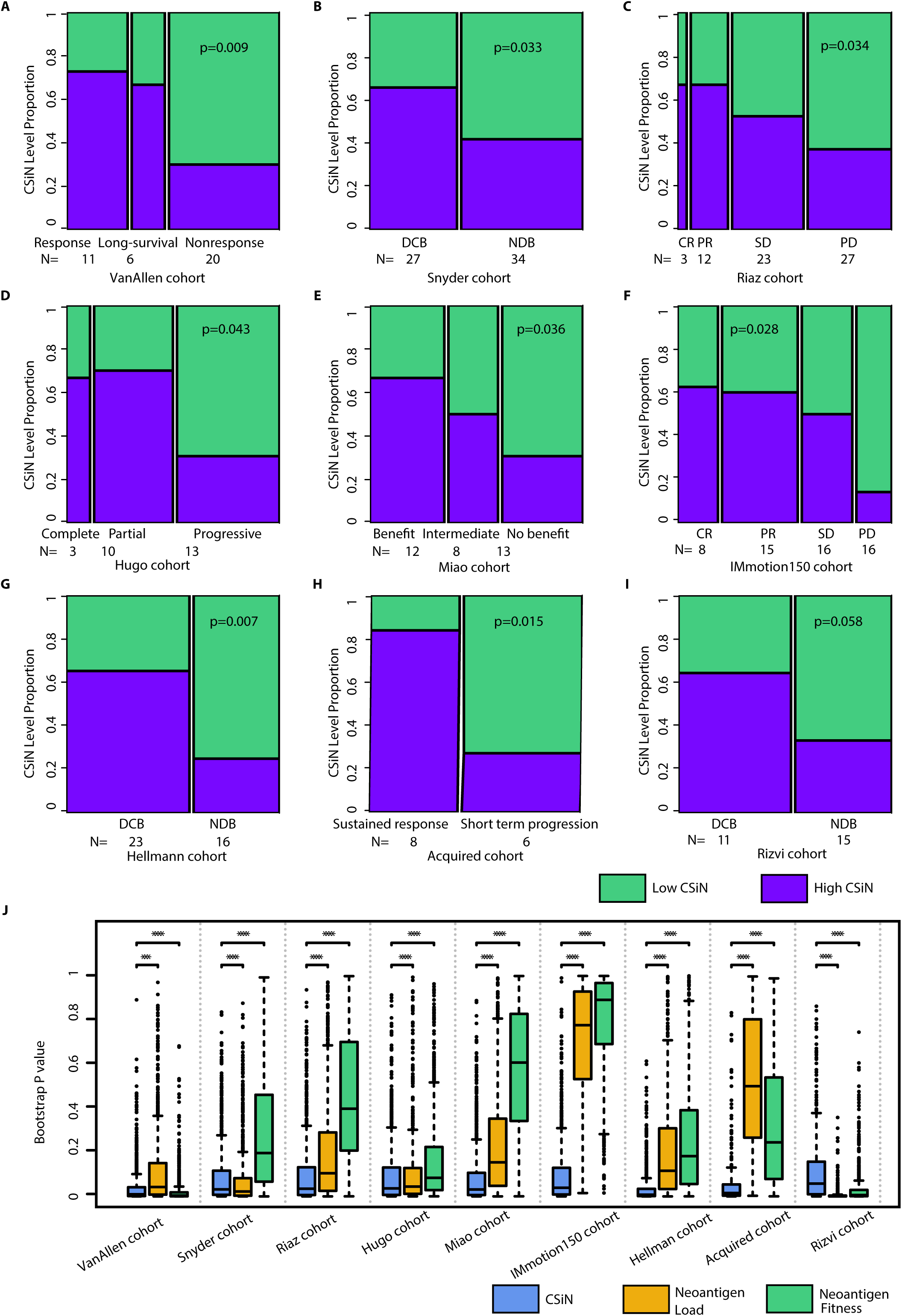
Association of CSiN with checkpoint inhibitor treatment response. (A) The VanAllen cohort. 11 patients with clinical benefit (response group), 6 patients with long-term survival with no clinical benefit (long-survival) group, and 20 patients with minimal or no clinical benefit (nonresponse) group. (B) The Snyder cohort. 27 patients with DCB, and 34 patients with NDB. (C) The Riaz cohort. 3 patients with complete response (CR), 12 patients with partial response (PR), 23 patients with stable disease (SD), and 27 patients with progressive disease (PD). (D) The Hugo cohort. 3 patients with complete response, 10 patients with partial response, and 13 patients with progressive disease. (E) The Miao cohort. 12 patients with clinical benefit, 8 patients with intermediate benefit, and 13 without clinical benefit. (F) The IMmotion150 cohort. There were 8 patients with CR, 15 patients with PR, 16 patients with SD, and 16 patients with PD. These patients were treated with atezolizumab and possess high T_eff_ signature expression. (G) The Hellmann cohort. There were 23 PD-L1+ (IHC>=3) patients with DCB, and 16 PD-L1+ patients with NDB. (H) The Acquired cohort. There were 8 patients with short term progression (progression<12 month) and 6 patients with sustained response (progression>12 month). (I) The Rizvi cohort. 11 patients with DCB and 15 patients with NCB. Biopsy and genomics data were obtained close to time of progression for all patients, while baseline biopsies were lacking for many patients. For (A)-(I), we tested the association of the dichotomized CSiN scores with the ordered response categories using an ordinal Chi-Square test. (J) Boxplots of bootstrap P values evaluating the robustness of the predictive performance of CSiN, neoantigen load and the neoantigen fitness score, with each P value generated from a bootstrap resample of each cohort. Two-sided Wilcoxon signed-rank test was used to compare the bootstrap P values. NS: P>0.01, *: P=0.01-0.05, **: P=0.001-0.01, ***: P=0.0001-0.001, ****:P<0.0001.

In comparison, we examined the predictive power of neoantigen load (**Fig. S1**) and the neoantigen fitness model (**Fig. S2**) in the same cohorts and by the same statistical tests. For consistency we used median to split the cohorts. For neoantigen load, we also adopted another cutoff (median + 2 x interquartile range), developed by Zehir *et al* (*38*) (results shown in **Supplementary Information**). For this analysis, the neoantigen fitness model is calculated for neoantigens of both class I and class II, as is done for CSiN and neoantigen load. We provided the results of the neoantigen fitness scores calculated using class I 9-mer neoantigens from missense mutations in **Supplementary Information**, as is done in its original report. Overall, neoantigen load and the neoantigen fitness model are not as strongly predictive as CSiN. We used bootstrap analysis to evaluate the statistical significance of the advance of CSiN against the other two approaches, which is an accepted methodology for model comparison (*39, 40*). In **Fig. 2J**, we showed that CSiN significantly outperformed neoantigen load in 7 out of the 9 cohorts evaluated and outperformed neoantigen fitness also in 7 out of the 9 cohorts. Overall, our results show that CSiN is capable of predicting response to checkpoint inhibitors in immunogenic cancers, and demonstrated a significant improvement over existing predictive tools. We also demonstrated the predictive power of CSiN, neoantigen load, and neoantigen fitness model using survival analysis criterion. Detailed results are shown in **Supplementary Information**.

Finally, we also explored the predictive power of CSiN in other forms of immunotherapies. We generated genomics data for an in-house cohort (the IL2 cohort) of ccRCC patients treated with concurrent IL-2 and Stereotactic Ablative Body Radiation (SAbR) treatment. IL2 has pleiotropic activating effects on cytotoxic T cells, T_reg_ cells and NK cells (*41*). It has been shown that SAbR has multiple immunogenic properties and could enhance the response to IL-2 (*42*). The neoantigen-mediated cytotoxicity probably partially explains the effects of this regimen. In this cohort, CSiN scores of patients with DCB are higher than CSiN scores of patients with NCB with marginal significance (**Fig. S3. A**, P=0.053), and out-perform neoantigen load and the neoantigen fitness model (**Fig. S3. B-C**).

### Higher CSiN predicts more favorable prognosis in immunogenic cancers

To understand the implication of neoantigen heterogeneity for long term survival of patients, we examined the association between CSiN and prognosis in the RCC, LUAD, LUSC and SKCM cohorts. We first focused on patients with high levels of T cell infiltration, profiled by our recently published eTME gene signatures (*31*). We speculated that the neoantigen-T cell axis is more likely to be functionally active when T cell infiltration is present in the tumor. Interestingly, in these patients, we indeed observed that higher CSiN scores had a significantly positive association with better survival for RCC (P=0.01, **Fig. 3A**), LUAD (P=0.036, **Fig. 3B**), LUSC (P=0.024, **Fig. 3C**), and SKCM (P=0.038, **Fig. 3D**). The False Discovery Rates for the 4 cohorts fall under 10% (**Sup. Table 3**). However, the overall survival of patients with lower T cell infiltration was indifferent to the levels of CSiN scores, which fits our speculation. We extracted and combined the high T cell infiltration patients from all four cohorts, and carried out survival analyses, which again showed that patients with higher CSiN scores had a significantly better overall prognosis (P=3.8×10^−5^, **Fig. 3E**). To further exclude the effect of clinical confounders, we performed multivariate survival analysis adjusted by disease type, stage, gender and age in this combined cohort. The significant association between survival and CSiN was retained (P<0.001, **Fig. 3F**).

**Fig. 3.**
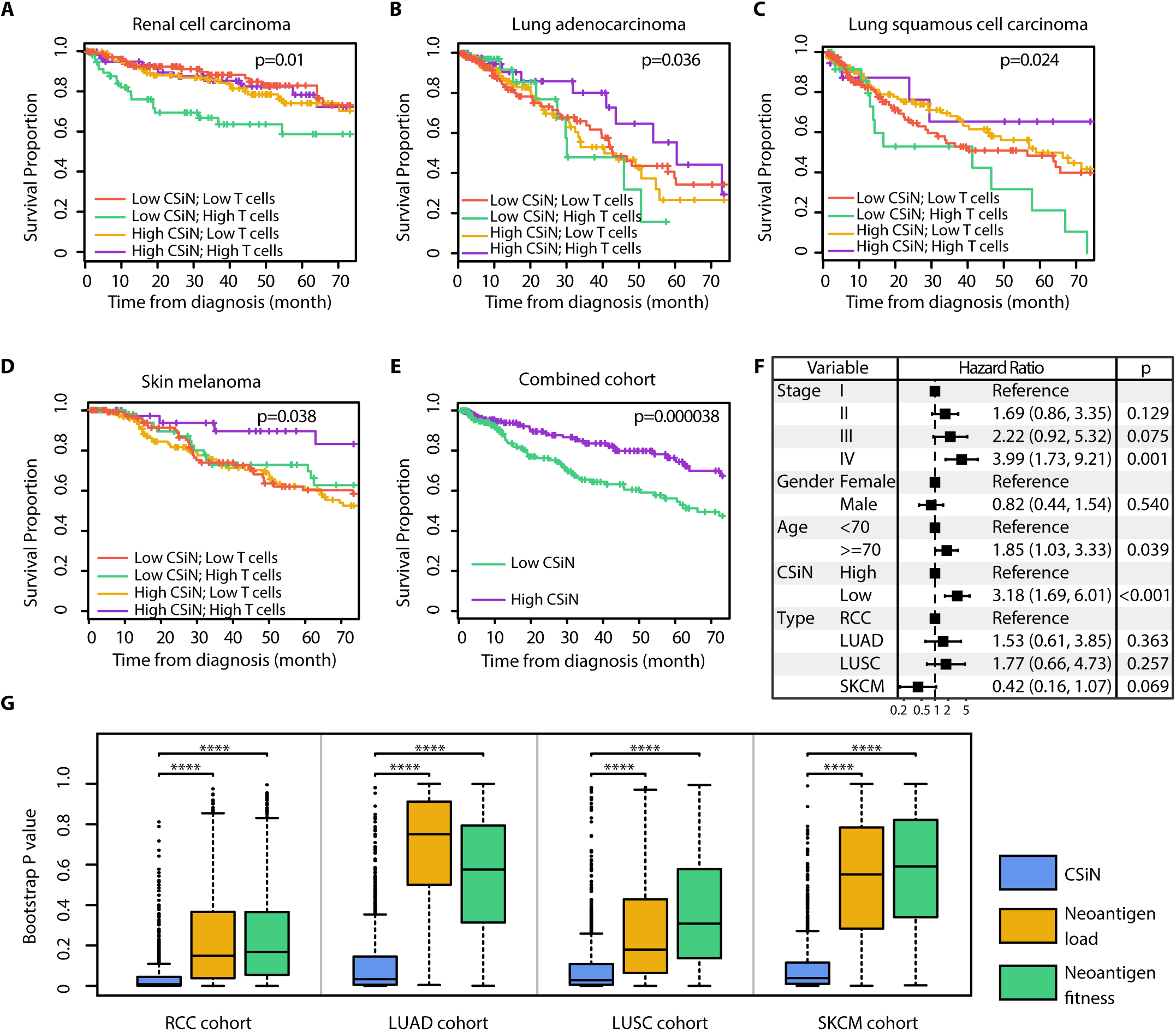
Association of CSiN with overall survival of patients. (A-E) Kaplan-Meier estimator was used to visualize patient overall survival. P values for logrank tests are shown. (A) The RCC cohort. (B) The LUAD cohort. (C) The LUSC cohort. (D) The SKCM cohort. (E) The patients identified as having “High T cells” are extracted from each cohort, combined, and tested together. The top 140 RCC patients, 100 LUAD patients, 100 SKCM patients, and 40 LUSC patients with the highest T cell infiltration were designated as having “High T cells”, according to their order of immunogenicity (*82*). The high and low CSiN score designations follow those in (A-D). (F) Forest plot for the coefficients of the multivariate CoxPH analysis of the combined cohort in (D). Disease type, pathological stage, gender, age and the binarized CSiN were included as covariates. The dotted line shows the no effect point. 95% Confidence intervals were shown as bars. (G) Boxplots of bootstrap P values evaluating the robustness of the prognostic performance of CSiN, neoantigen load and the neoantigen fitness score, with each P value generated from a bootstrap resample of each cohort. Two-sided Wilcoxon signed-rank test was used to compare the bootstrap P values. *: P=0.01-0.05, **: P=0.001-0.01, ***: P=0.0001-0.001, ****:P<0.0001.

In contrast, the same analysis for neoantigen load and the neoantigen fitness model yielded insignificant association (**Fig. S4** and **Fig. S5**). We also employed the bootstrap analysis to evaluate the statistical significance of this comparison. In **Fig. 3G**, we showed that CSiN significantly outperformed both methods in all 4 cohorts evaluated. Overall, in concordance with several previous studies that reported a lack of association of higher neoantigen load with better prognosis in several cancer types (*19*–*21*), our results suggest that the heterogeneity of neoantigens could be more prognostically important.

We assessed non-immunogenic cancer types as well. Pediatric Acute Lymphocytic Leukemia is an aggressive childhood tumor type with low neoantigen load. We evaluated a cohort (pALL) of pediatric ALL patients and observed that CSiN was not predictive of prognosis (**Fig. S6. A**, P=0.584), with the results for neoantigen load shown in **Fig. S6. B** and neoantigen fitness shown **in Fig. S6. C**). We considered the liver hepatocellular carcinoma patients from The Cancer Genome Atlas (TCGA) (LIHC cohort), although whether liver cancer is an immunogenic cancer type is still under debate (*43, 44*). We observed patients with higher CSiN scores had a non-significant trend of better survival than patients with low CSiN scores in the high T cell infiltration subset of patients (**Fig. S6. D**, P=0.165), with the results for neoantigen load shown in **Fig. S6. E** and neoantigen fitness shown in **Fig. S6. F**). These observations further confirmed the coupling effect of CSiN with the immunogenic environment of tumors.

## Discussion

The fundamentally surprising biological insight of our work is that the neoantigen clonal structure in each tumor specimen and the immunogenicity quality of neoantigens (represented by the MHC-binding strength in our study) are predictive of response to checkpoint inhibitors and prognosis. And this could significantly out-weight the simple neoantigen count. Our comprehensive analyses show that CSiN, which describes these properties of the neoantigen profile quantitatively, possesses substantially better predictive and prognostic performance than other neoantigen-based biomarkers, in the majority of evaluated cohorts. Our implementations of CSiN, neoantigen load, and neoantigen fitness model have considered both MHC class I and class II neoantigens, and also neoantigens generated from insertions/deletions and stop-loss mutations. This is different from the original publication of the neoantigen fitness model that only considered 9-mer class I neoantigens generated from missense mutations. We believe inclusion of all these sources of neoantigens is important for a complete characterization of the neoantigen profiles in each patient (analyses for each class of neoantigens only are shown in **Supplementary Information**). In alignment with our reports, McGranahan *et al* made a qualitative observation that CTLA-4-resistant tumors could be enriched for subclonal mutations, which may enhance total neoantigen burden but not elicit an effective antitumor response due to the subclonal nature of these neoantigens (*13*). Miao *et al* also made a similar observation (*25*). Our study is distinguished from these earlier reports in that we provided a robust quantitative measurement that was subjected to systematic evaluations, and we also evaluated prognosis in addition to treatment response. Overall, CSiN likely constitutes an important predictive tool for medical oncologists treating patients with checkpoint blockade, and has addressed the limitations of prior neoantigen-based predictive biomarkers of checkpoint inhibitors.

CSiN extracts genetics information that is not captured by neoantigen load, neoantigen fitness model or expression-based biomarkers. A number of expression-based biomarkers for immunotherapy have been proposed and validated on different levels so far. For example, PD-L1 expression in the tumor microenvironment (*45, 46*), Th1-type chemokine expression (*47*), and T cell infiltration (*48*), as well as many others. But CSiN augments and complements, rather than replaces, these biomarkers. We show that CSiN is associated with longer survival only in patients with sufficient T cell infiltration. Similarly in the treatment cohorts, the P values for testing the correlation between CSiN and treatment response in the high T cell/high PD-L1 patient subsets are generally smaller compared with the P values of the full cohorts (**Fig. 2** and **Supplementary Information**), even though the subset cohorts are much smaller in sample size.

These results suggest that it is crucial for all components of the neoantigen-host immune axis to be functionally active, in order to enable efficient immuno-elimination of tumor cells. The insignificant association of CSiN with prognosis in the less immunogenic liver cancers and pediatric ALLs also supports this notion. Our observations may inspire potential future studies to construct more sophisticated predictive and prognostic models that incorporate CSiN, neoantigen load, the neoantigen fitness model and other biomarkers together for improved performance.

Our results reporting the positive correlation between neoantigens and treatment response in RCCs is interesting. Currently, the field is still debating the role of neoantigens in immune response of RCCs. While RCCs have low neoantigen/mutation loads, Turajlic *et al* discovered that RCCs have the highest number of insertion/deletion mutations on a pan-cancer basis (*49*), which tend to encode high quality neoantigens. In terms of predicting survival after immunotherapy treatment, Samstein *et al* reported a significant correlation between tumor mutation burden and progression free survival (*50*), while this observation is not made in the phase 3 JAVELIN Renal 101 trial (*51*). Cherkasova *et al* discovered the re-activation of a HERV-E retrovirus in RCCs, which can encode an immunogenic peptide recognizable by cytotoxic T cells (*52*). It is highly likely both neoantigens and self antigens contribute to the immunogenicity in RCCs. In the future, it will be of interest to investigate whether CSiN is more predictive of patients’ response to immunotherapies when such antigens have been incorporated in the calculation, in more and larger cohorts.

One limitation of our study and, to some extent, this field, is that we used predicted neoantigens from genomics data for correlation with patient phenotypes. Despite our efforts to validate the neoantigen predictions, it is likely there are still false positive and false negative predicted neoantigens that convoluted our analyses. In future studies, incorporating the genomics-based approach with other methods, such as mass spectrometry(*53*), may improve the sensitivity and specificity of neoantigen detection, and thus further enhance the predictive power of CSiN.

Class I and class II neoantigens represent very different aspects of immune response. But several recent studies have implicated neoantigen-specific CD4^+^ T-cells in direct tumor clearance (*54*–*56*). Our results also suggest that the inclusion of class II neoantigens is important for treatment response prediction. In the future, it will be interesting to carry out comprehensive research into the roles of class II neoantigens in the tumor microenvironment.

The neoantigen repertoire is in a constant dynamic evolution (*2, 57*), in which immunoediting and immunotherapy treatment actively modify its landscape. CSiN offers a new tool to monitor the neoantigen profiles, where different tumor cells could have different growth advantages subject to the pressure of T cell cytotoxicity determined by each cell’s neoantigen composition. Overall, our work offers a rigorous methodology of predicting response to immunotherapy and prognosis from routine patient samples, and should be useful for personalizing medicine in the modern era of immunotherapy.

## Materials and Methods

### Study Design

The objective of this research project is to study the implication of the clonal structure of neoantigens for predicting treatment response and prognosis. The research subjects are individual cancer patients. This is a descriptive study. Patients were or were not treated with immunotherapy. As this is a retrospective analysis study, the researchers were not blinded to the allocation labels. We included all available samples and data from either public or private sources into our study. We stopped the data collection on the lock date of September 1st, 2019. The endpoints considered are response categories and survival of cancer patients treated with immunotherapy or baseline cancer patients. Usually one sample per patients was available. In the uncommon cases of more than one sample collected for each patient, we will average the statistics (*e*.*g*. CSiN) calculated for each patient.

### The Cauchy–Schwarz index of Neoantigens

In CSiN, we considered the pairing between the repertoire of neoantigens and the tumor mutations to which they belong. One way to characterize this property is to average the product of the variant allele frequencies (VAF) of somatic mutations and the number of neoantigens generated by each mutation, normalized by the average VAF and average mutation-specific neoantigen load in each patient. It is >1 under a “good” pairing, and *vice versa*. This forms the backbone of the final CSiN score. CSiN was so named as the pairing of tumor mutations and neoantigens and its effect on the overall CSiN score bear analogy to the Cauchy-Schwarz inequality, which describes the upper bound of the product sum of two vectors of real numbers and the condition for the equality to be achieved - the values of the two vectors are in parallel (naturally then in the same ranked order). Refer to **Supplementary Information** for the implementation details of CSiN and additional analyses involving CSiN, which are not shown in the main text. A cartoon showing the workflow of CSiN is shown in **Fig. S7**.

### The QBRC mutation calling pipeline

We used the QBRC mutation calling pipeline for somatic mutation calling, developed in the Quantitative Biomedical Research Center (QBRC) of UT Southwestern Medical Center. In short, exome-seq reads were aligned to the human reference genome by BWA-MEM (*58*). Picard was used to add read group information and sambamba was used to mark PCR duplicates. The GATK toolkit (*59*–*61*) was used to perform base quality score recalibration and local realignment around Indels. MuTect (*62*), VarScan (*63*), Shimmer, SpeedSeq (*64*), Manta, and Strelka2 (*65*) were used to call SNPs and Indels. A mutation that was repeatedly called by any two of these software was retained. Annovar was used to annotate SNPs and Indels, and protein sequence changes (*66*). All SNPs and Indels were combined and only kept if there were at least 7 total (wild type and variant) reads in the normal sample and at least 3 variant reads in the tumor sample. Somatic mutations and germline mutations were annotated according to the variant allele frequencies in the normal and tumor samples.

### The QBRC neoantigen calling pipeline

We used the QBRC neoantigen calling pipeline for neoantigen calling, which follows similar standards as those pipelines used in high impact publications such as (*17, 18, 22, 34, 67, 68*). The validity of our neoantigen predictions is demonstrated in **Supplementary Information**. We kept only frameshift, non-frameshift, missense and stop-loss mutations that would lead to protein sequence changes. We kept only somatic mutations whose variant allele frequencies (VAFs) were <0.02 in the normal sample and VAFs>0.05 in the tumor samples. For class I HLA proteins (A, B, C), we predicted putative neoantigens of 8-11 amino acids in length, and for class II HLA proteins (DRB1 and DQB1/DQA1), we predicted putative neoantigens of 15 amino acids in length. Class I and II HLA subtypes were predicted by the ATHLATES tool (*69*), which has been shown to be accurate for these HLA alleles (*70*). Samples whose total successfully typed HLA alleles (counting both chromosomes) were <8 were regarded as poor quality data, and were left out from downstream analyses. Putative neoantigens with amino acid sequences exactly matching known human protein sequences were filtered out. For class I bindings, the IEDB-recommended mode (http://tools.iedb.org/main/) was used for prediction of binding affinities, while for class II binding, NetMHCIIpan embedded in the IEDB toolkit was. Neoantigens were kept only if the predicted ranks of binding affinities were ≤2%. Tumor RNA-seq data, if available, were aligned to the reference genome using the STAR aligner (*71*). FeatureCounts was used to summarize gene expression levels (*72*). Neoantigens whose corresponding mutations were in genes with expression level <1 RPKM in either the specific exon or the whole transcript were filtered out. For the samples analyzed by our pipeline, we applied the filtering by RNA-seq data on the neoantigen list when RNA-Seq data are available. We showed the results for calculating CSiN with only exome-Seq data in **Supplementary Information**, which indicated that filtering the neoantigen list by RNA-Seq data is important for the predictive performance of CSiN. Finally, in accordance with our pipeline, evidence from Ott *et al* (*73*) has shown that neoantigens of class I and class II, and also different registers can all possibly be immunogenic, and IEDB (https://www.iedb.org/) has documented many immunogenic antigens of different peptide lengths.

### Patient cohorts

For all patient cohorts, if approval of access to raw exome-seq and RNA-Seq data had been obtained, we predicted the somatic mutations and neoantigens using our in-house pipelines. In cases where raw genomics data were not accessible, we analyzed the processed neoantigen and mutation data, which were included in the supplemental files of the original publications. For exploratory analysis and association with overall survival, we collected data from 110 RCC patients from the Kidney Cancer Program at UT Southwestern Medical Center (*31*), 94 ccRCC patients from Sato *et al (23)*, and 162 ccRCC patients from TCGA (*74*) (the RCC cohort); 427 lung adenocarcinoma patients from TCGA (*75*) (the LUAD cohort); 389 lung squamous cell carcinoma patients from TCGA (*76*) (the LUSC cohort); 401 melanoma patients from TCGA (*77*) (the SKCM cohort); 103 pediatric and young adult T-lineage acute lymphoblastic leukemia patients (the pALL cohort) from Liu *et al (24)*; and 292 liver cancer patients from TCGA (*78*) (the LIHC cohort) (**Sup. Table 1**).

For association of CSiN scores with immunotherapy response, 10 patient cohorts were collected and summarized in **Sup. Table 1. Sup. Table 1** shows the total number of patients for each cohort used in correlation of treatment response and neoantigen-based predictive markers in this study. In the VanAllen cohort, there are 40 patients with matched RNA-Seq and exome-seq data and 3 patients were further removed due to lack of response information. For the Riaz cohort, 3 patients were removed as two of them have lack of response information (“not evaluated” reported by the original study) and the third patient’s HLA alleles cannot be typed accurately from the sequencing data. In the Miao cohort, 2 patients were removed due to the HLA typing issue. In the Rizvi cohort, 3 patients were removed due to lack of response information (not reaching 6 month follow up) and 5 patients were removed due to the HLA typing issue. In the Snyder cohort, 3 patients were removed as the neoantigen data were not available from the original publication (we used neoantigens called by the original study for this cohort as we don’t have access to the raw genomics data). In the IMmotion150 cohort, 99 patients on atezolizumab and 50 patients on sunitinib have both exome-seq and RNA-Seq. 3 patients treated by atezolizumab and 4 patients treated by sunitinib were further removed because response information is not available. For the Hellmann cohort, the neoantigen lists were generated by the original authors and used by us in this work. There are 74 patients with neoantigen data generated. Only whole exome-seq was done and only about half of these patients have consented to genomics data sharing, so we decided to use these neoantigens called by the original report. For the Hugo cohort, 28 have both exome-seq and RNA-Seq data, and 2 were further removed due to the HLA typing issue. Stratified analyses (**Fig. 2** and **Supplementary Information**) were performed for the IMmotion150, VanAllen, Hugo, Riaz and Miao cohorts (with RNA-Seq data available) based on ssGSEA analyses (*79*) of the T_eff_ gene signature (*80*) to focus on the top 60% patients, and also performed for the Hellmann cohort based on PD-L1 IHC level with a cutoff on IHC readings (which are given in integers by the original report for 69 patients) chosen such that the top subset is closest to 60% of the whole cohort. For cases in which multiple types of treatment outcomes were recorded in a cohort, we used the primary criterion employed by the original publications. Patient samples that were analyzed but were not shown in **Fig. 2** (*e*.*g*. IMmotion150 patients on the sunitinib arm) were shown in **Supplementary Information**.

### Statistical analyses

All computations and statistical analyses were carried out in the R computing environment. For all boxplots appearing in this study, box boundaries represent interquartile ranges, whiskers extend to the most extreme data point which is no more than 1.5 times the interquartile range, and the line in the middle of the box represents the median. For association of CSiN, neoantigen load and the fitness model with binary/categorical response variables, we tested the association of the dichotomized CSiN scores with the ordered response categories (*e*.*g*. CR->PR->SD->PD) using an ordinal Chi-Square test. The three types of predictive scores are each split on the median within each cohort, to ensure comparability and avoid the scaling problem. The alternative hypothesis is that patients with better response and survival outcome are more likely to have higher CSiN scores. We employed the *chisq_test* function in the R *coin* package for this purpose. For survival-type analysis, we adopted the log-rank test for evaluating whether patients with higher CSiN scores had better prognosis. T cell infiltrations and activation of the IFN-gamma signature were predicted using the single sample gene set enrichment analysis (ssGSEA) method (*79*). ssGSEA analysis was performed using the R *GSVA* package by calling the *gsva* function with parameter method="ssgsea” and rnaseq=T (*81*). The forest plot was performed by the *forest_model* function in the R *forestmodel* package. For model comparison, 5,000 bootstrap resamples of each cohort were generated, and each resample was used to evaluate the predictive or prognostic performance of CSiN, neoantigen load and the neoantigen fitness score. The P values of 5,000 bootstraps of each approach were compared using two-sided Wilcoxon signed-rank test.

## Supplementary Materials

**Fig. S1** Predictive power of neoantigen load.

**Fig. S2** Predictive power of the neoantigen fitness model.

**Fig. S3** Association of CSiN (a), neoantigen loads (b) and neoantigen fitness (c) with IL2/SAbR treatment response in ccRCC patients.

**Fig. S4** Prognostic power of neoantigen load.

**Fig. S5** Prognostic power of the neoantigen fitness model.

**Fig. S6** Association of CSiN, neoantigen loads and neoantigen fitness with prognosis of pediatric ALL patients and LIHC patients.

**Fig. S7** Cartoon showing the workflow of calculation of CSiN scores.

**Sup. Table 1** The patient cohorts used in this study.

**Sup. Table 2** Processed mutation, expression and neoantigen data of the IL2 cohort

**Sup. Table 3** P values and False Discovery Rates of the tested cohorts shown in **Fig. 2** and **Fig. 3**

**Sup. Table 4** Raw Data Table of some data used in **Fig. 1, Fig. 2**, and **Fig. 3**

**Supplementary Information** A comprehensive explanation of the CSiN score and additional characterization of CSiN score.

## Acknowledgements

We acknowledge Jessie Norris for her helpful comments on the writing of the paper. We acknowledge Dr. Ying Wang for her helpful comments on the scientific contents. We acknowledge the Wakeland sequencing core of UT Southwestern for generating genomics data for the IL2 cohort. We acknowledge the authors of the phs000452.v2.p1, the phs001493.v1.p1, the phs001464.v1.p1, the phs000218.v1.p1, and the phs000464.v15.p7 datasets, as well as the funding agencies that supported these studies and dbGaP that supported the archival of these datasets.

## Author contributions

T.L., S.W., Q.Z., and Z.X. contributed to computational analyses. T.W. contributed to the overall supervision of the project. L.X., and J.B. contributed to data access. S.M., L.P., M.C., and R.H. contributed to the collection of samples and generation of sequencing data for the IL2 cohort. L.X., N.S., J.G., R.H., J.B., and T.W. contributed to study design and manuscript writing.

## Data and material availability

The QBRC mutation calling and neoantigen calling pipelines are available on GitHub: https://github.com/tianshilu/QBRC-Somatic-Pipeline and https://github.com/tianshilu/QBRC-Neoantigen-Pipeline. The CSiN calculations were embedded in the neoantigen pipeline. **Sup. Table 1** lists the data source and/or accession methods of these studies. The raw RNA-Seq and exome-seq data of the IL2 cohort patients can be downloaded from the European Genome phenome Archive with accession number EGAS00001003605 through controlled access. The processed mutation and neoantigen data of the IL2 cohort can be found in **Sup. Table 2**.

## Supplementary Materials

**Fig. S1** Predictive power of neoantigen load. The analyses are the same as in **Fig. 2**, except that neoantigen loads are considered.

**Fig. S2** Predictive power of the neoantigen fitness model. The analyses are the same as in **Fig. 2**, except that neoantigen fitness model is considered.

**Fig. S3** Association of CSiN (A), neoantigen loads (B) and neoantigen fitness (C) with IL2/SAbR treatment response in ccRCC patients. 3 patients with complete response (CR), 1 patient with partial response (PR), and 2 patients with stable disease (SD) for more than 6 months form the DCB group. 3 patients with stable disease (SD) less than 6 months and 7 patients with progressive disease (PD) form the NCB group.

**Fig. S4** Prognostic power of neoantigen load. The analyses are the same as in **Fig. 3**, except that neoantigen loads are considered.

**Fig. S5** Prognostic power of the neoantigen fitness model. The analyses are the same as in **Fig. 3**, except that neoantigen fitness model is considered.

**Fig. S6** Association of CSiN, neoantigen loads and neoantigen fitness with prognosis of pediatric ALL patients and LIHC patients. P values for logrank tests are shown. (A-C) 103 pediatric and young adult T-lineage acute lymphoblastic leukemia patients were analyzed. (D-F) 292 TCGA LIHC patients were analyzed. The top 40 LIHC patients were designated as having “High T cells”, as LIHC is less immunogenic than the other tumor types investigated in this study.

**Fig. S7** Cartoon showing the workflow of calculation of CSiN scores.

**Sup. Table 1** The patient cohorts used in this study.

**Sup. Table 2** Processed mutation, expression and neoantigen data of the IL2 cohort

