## Supplementary figures and images for "Tumor Neoantigenicity Assessment with CSiN Score Incorporates Clonality and Immunogenicity to Predict Immunotherapy Outcomes"

### Sup. Fig. 1

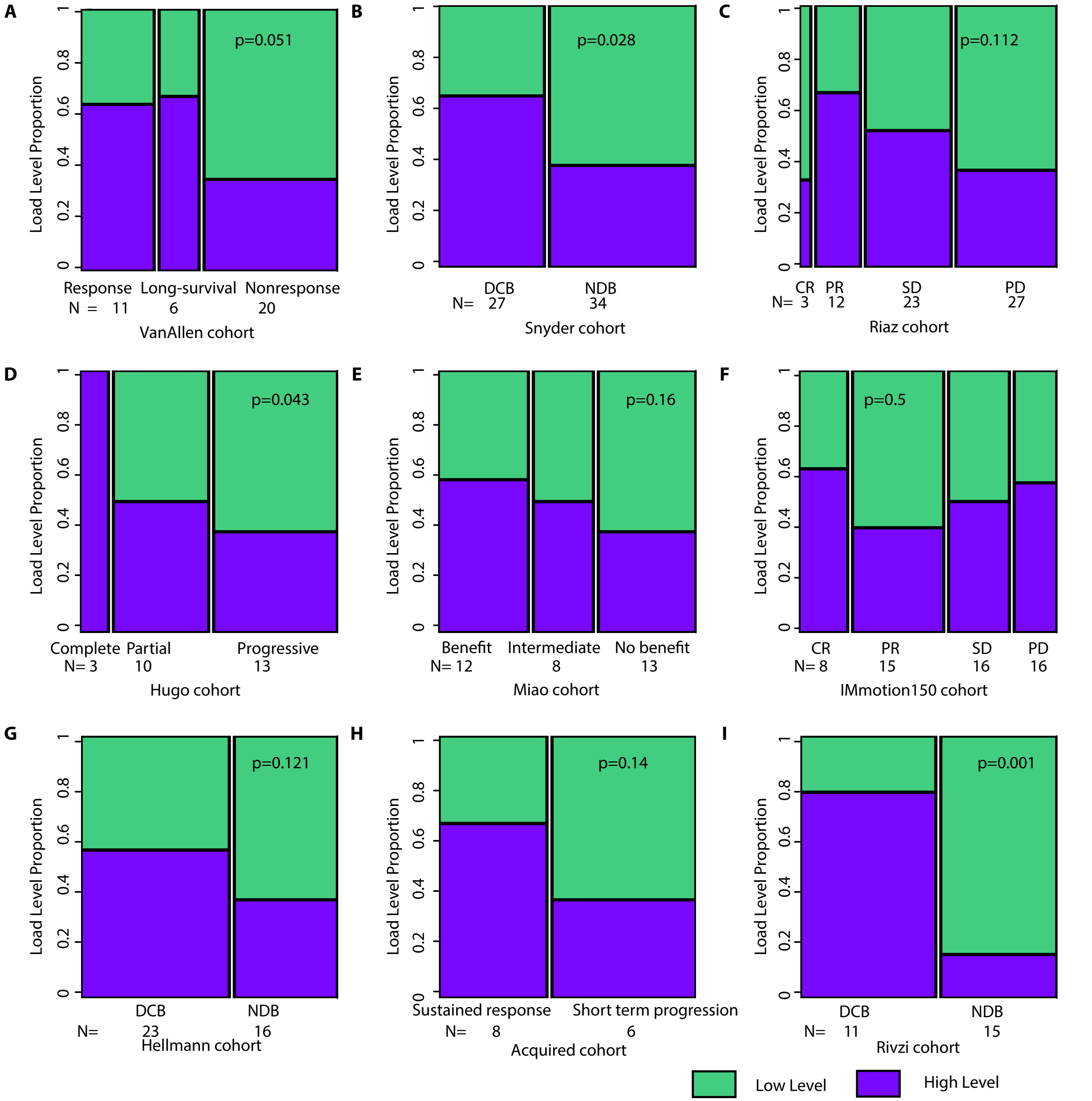

### Sup. Fig. 2

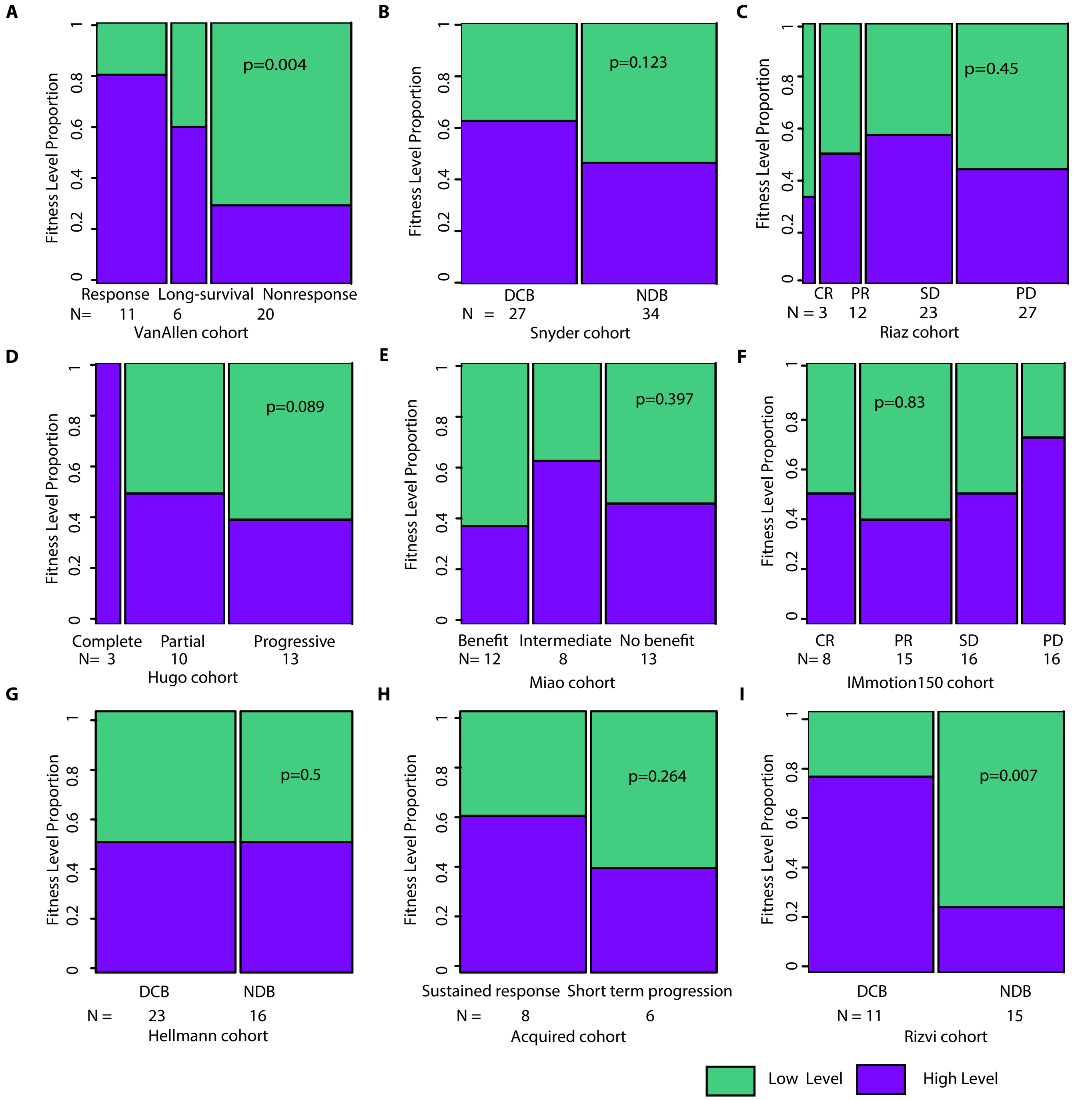

### Sup. Fig. 3

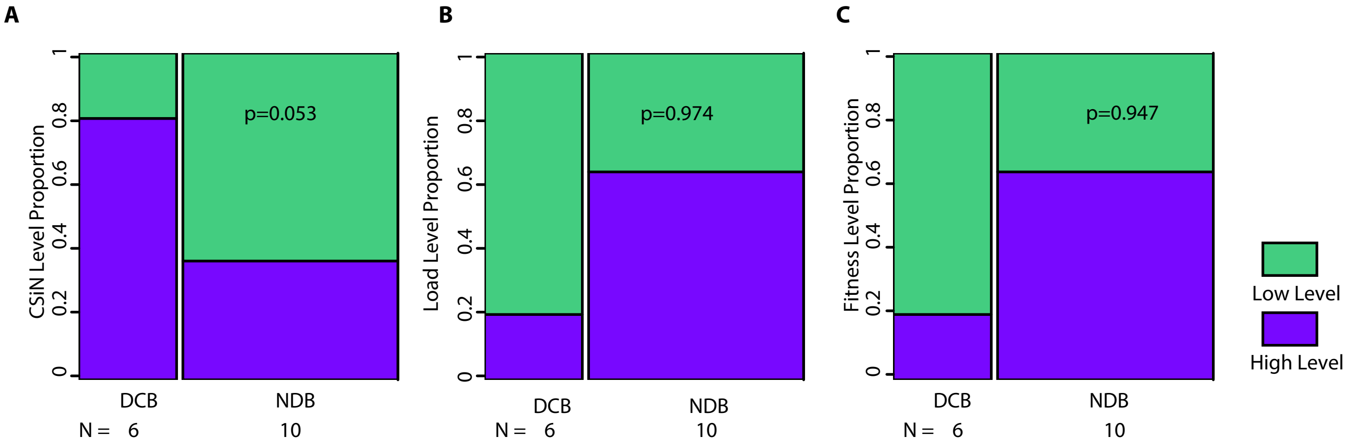

### Sup. Fig. 4

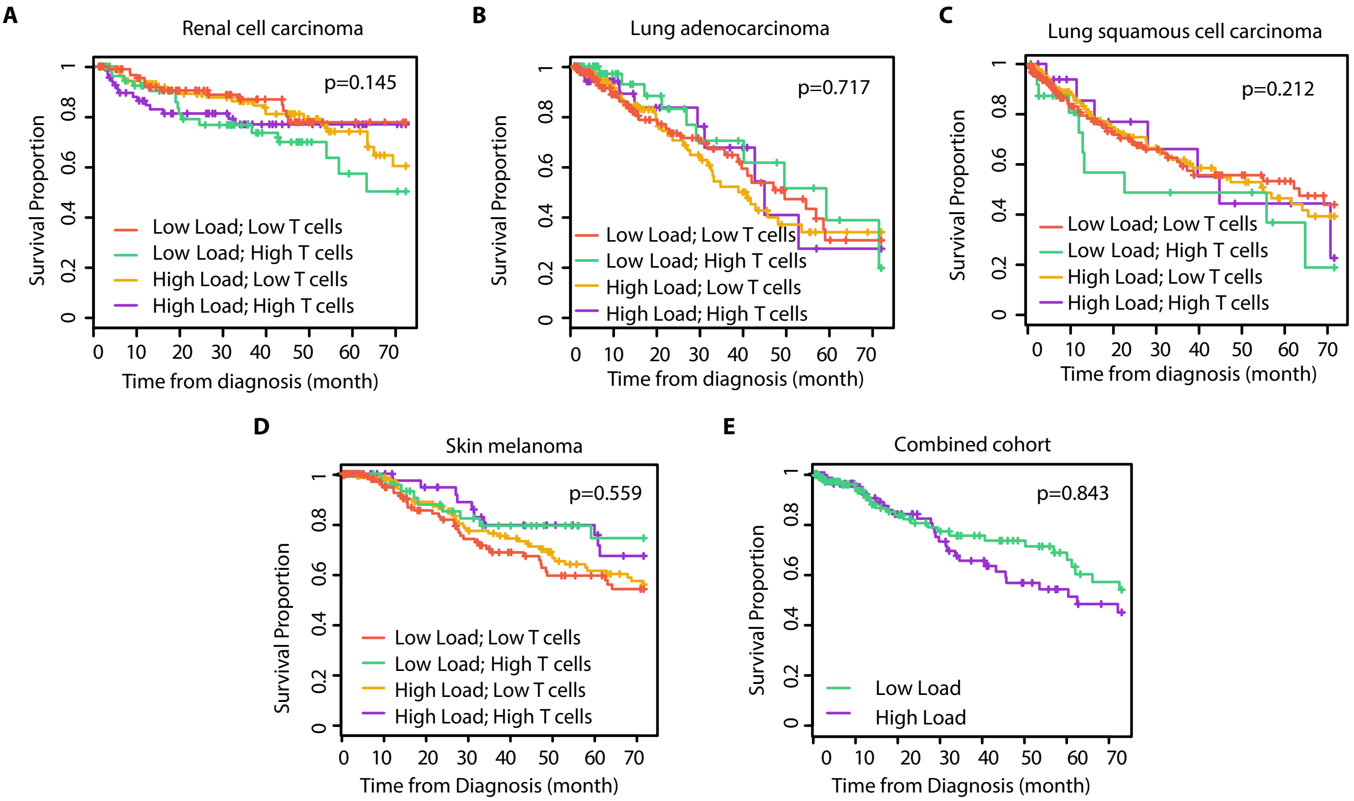

### Sup. Fig. 5

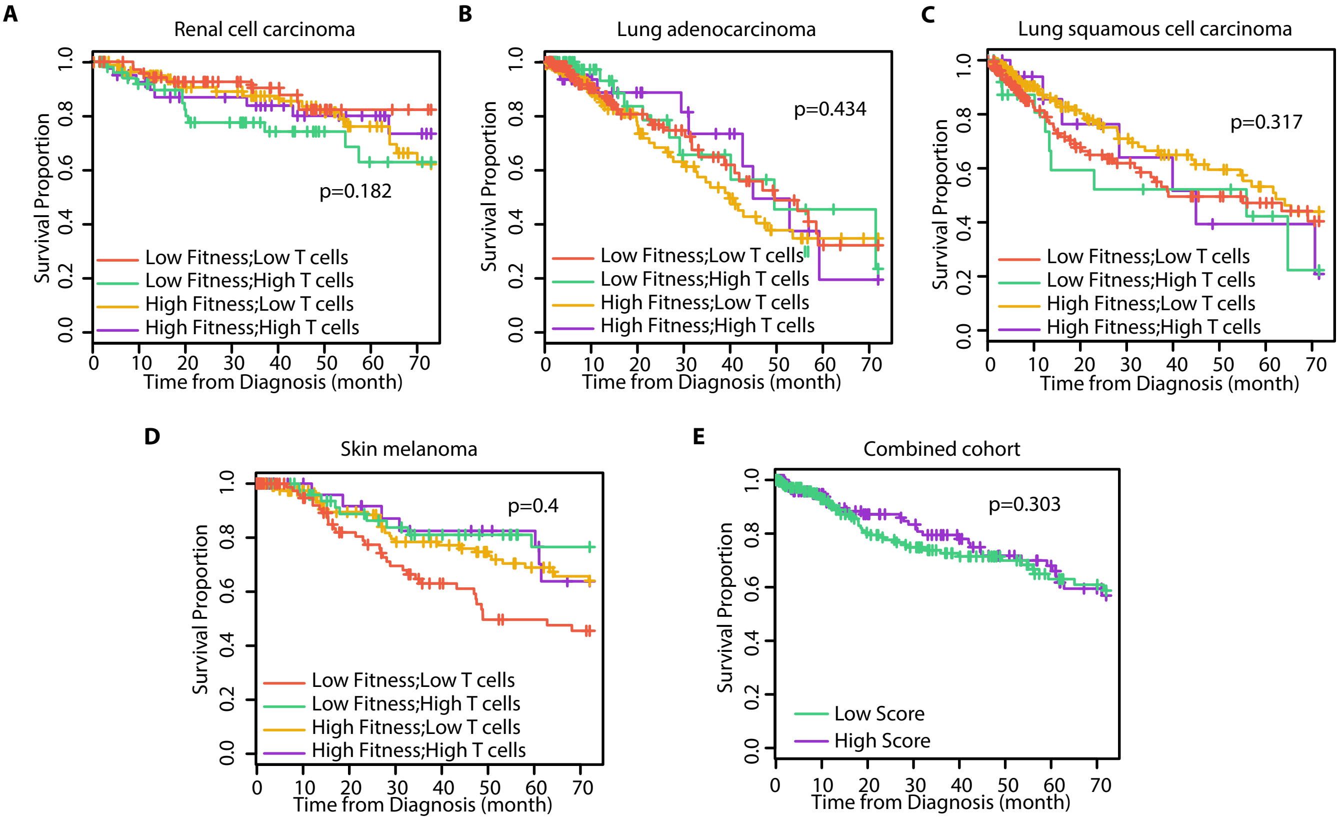

### Sup. Fig. 6

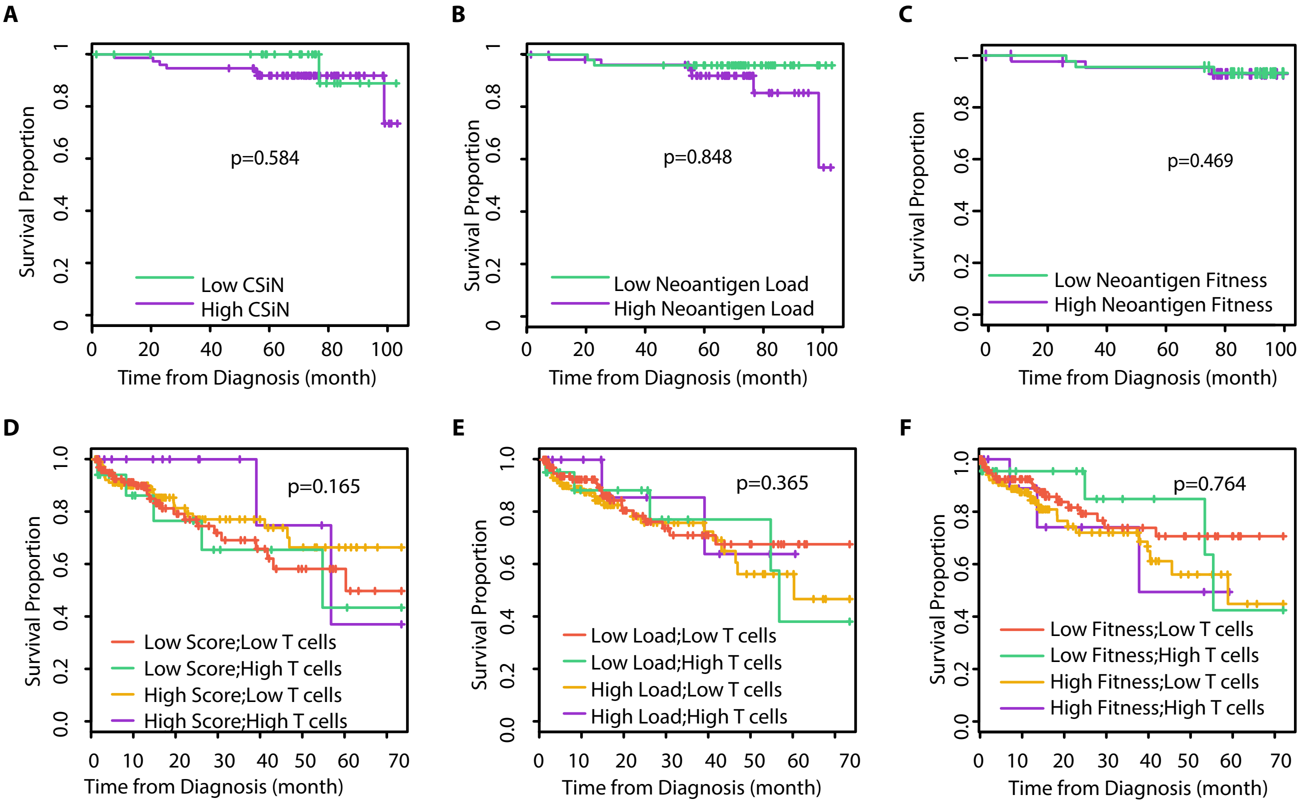

### Sup. Fig. 7

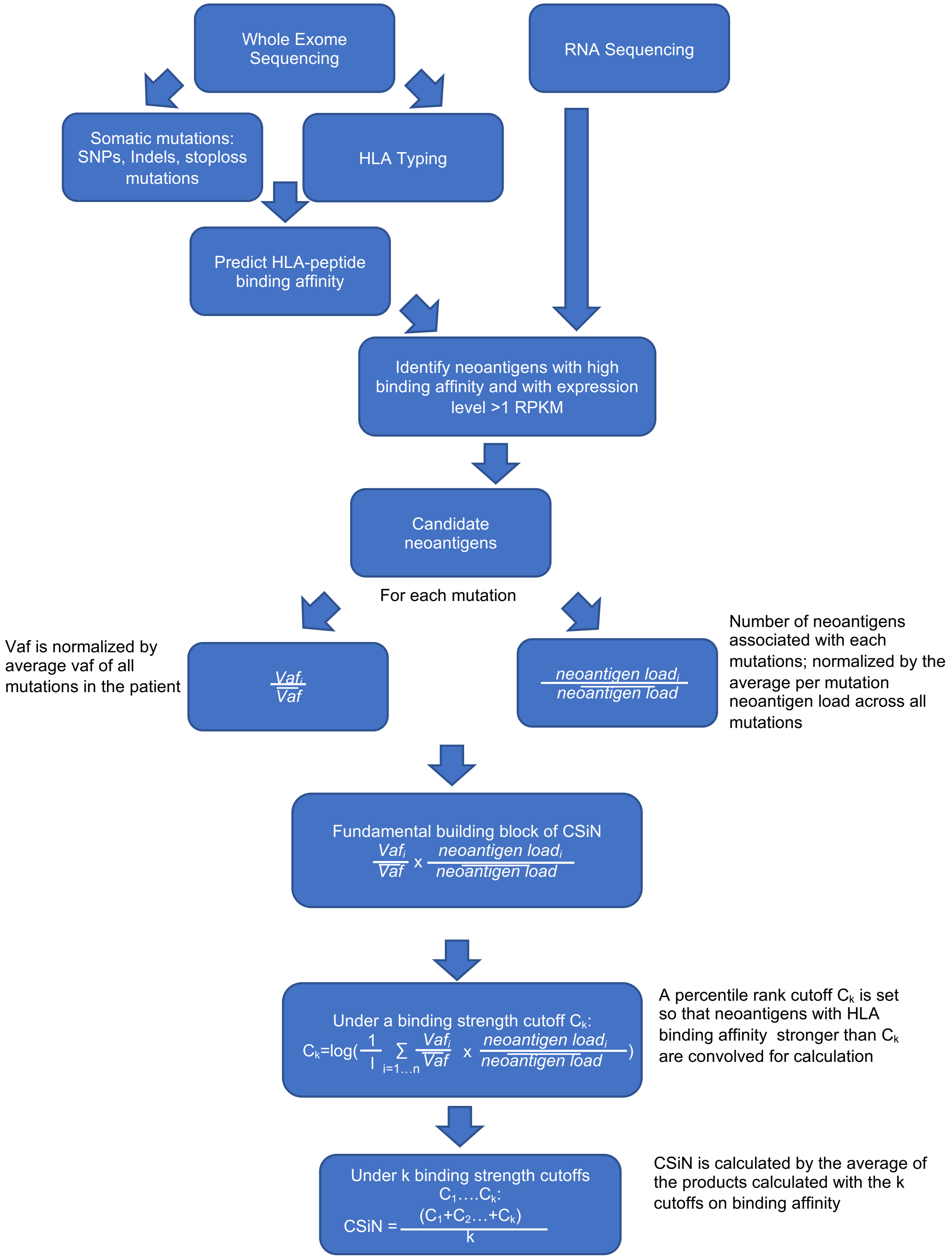
