## Supplementary Information for "Tumor Neoantigenicity Assessment with CSiN Score Incorporates Clonality and Immunogenicity to Predict Immunotherapy Outcomes"

### Detailed explanation of CSiN

**Definition:**

(1) The fundamental building block of CSiN is $\sum_{i=1..n} \frac{Vaf_i}{\overline{Vaf}} \times \frac{load_i}{\overline{load}}$. The variance allele frequency (VAF) is the number of variant reads divided by the total number of reads covering reach variant position. The load is the number of neoantigens associated with each mutation. $n$ is the total number of missense, indels, and stoploss somatic mutations in a tumor sample. $\overline{Vaf}$ describes the average VAF of all the somatic mutations (to control for tumor purity) and $\overline{load}$ is the average per mutation neoantigen load across all somatic mutations (so CSiN is orthogonal to neoantigen load). It is common to see different tumor biopsies have different levels of non-tumor cell contents (immune and stromal cells), and the tumor mutations' VAFs will be influenced by this confounding factor. The procedure of division by $\overline{Vaf}$ helps to normalize this effect.

According to the Cauchy-Schwarz inequality, when the mutations with higher VAFs are also the mutations that generate more neoantigens (our hypothesized favorable distribution), the product value will be larger (higher CSiN score). Therefore, a higher CSiN conforms to a favorable neoantigen clonal structure.

(2) Because the neoantigens vary in quality, and to give more weight to better neoantigens, the value is calculated by the average of the products calculated with different cutoffs on quality of neoantigens, with better neoantigens convolved in more rounds of calculations.

$$CSiN = \frac{\sum_{c=\{c_0,c_1,\ldots c_k\}} \log\left(\frac{\sum_{i=1..n,\ q(i)>c} \frac{Vaf_i}{\overline{Vaf_c}} \times \frac{load_i}{\overline{load_c}}}{\sum_{i=1..n} I(q(i)>c)}\right)}{k}$$

In this study, we used the percentile rank variable generated by the IEDB MHC binding affinity prediction software as the quality metric, $q(i)$. This variable measures the binding strength between neoantigens and the MHC molecules, and a smaller percentile rank delineates a greater affinity. The average VAF and neoantigens load are calculated with their according cutoff value, $c$, and we used $k$ cutoff values of *0.375, 0.5, 0.625, 0.75, 1.25, 1.75,* and *2*. The upper bound of the cutoff values is 2%, which is the most well established cutoff for an epitope to be considered as an HLA binder, according to netMHCpan. $I(s)$ evaluates to 1 if the statement $s$ is true, 0 otherwise. Accordingly, the definition of the average VAF and neoantigen loads are revised as:

$$\overline{Vaf_c} = \frac{\sum_{\substack{i=1..n \\ q(i)>c}} Vaf_i}{\sum_{i=1..n} I(q(i)>c)} \text{ and } \overline{load_c} = \frac{\sum_{\substack{i=1..n \\ q(i)>c}} load_i}{\sum_{i=1..n} I(q(i)>c)}$$

(3)

$$CSiN = \frac{\sum_{c=\{c_0,c_1,...c_k\}} \log\left(\frac{\sum_{\substack{i=1..n \\ rank(-Vaf_i)\le M \\ q(i)>c}} \frac{Vaf_i}{\overline{Vaf_c}} \times \frac{load_i}{\overline{load_c}}}{\sum_{\substack{i=1..n \\ rank(-Vaf_i)\le M}} I(q(i)>c)}\right)}{k}$$

To accommodate the patient samples with extremely large number of mutations, an adjustment is made where the calculation only considers the top M mutations with the largest VAFs when there are more than M mutations (M=500 in this study).

(4) The CSiN score defined above is a random variable centered approximately at zero. The final reported CSiN score is multiplied by a fixed constant, a (a=10), to increase the dynamic range for better visualization.

$$CSiN = \frac{a}{k} \times \sum_{c=\{c_0,c_1,...c_k\}} \log\left(\frac{\sum_{\substack{i=1..n \\ rank(-Vaf_i)\le M \\ q(i)>c}} \frac{Vaf_i}{\overline{Vaf_c}} \times \frac{load_i}{\overline{load_c}}}{\sum_{\substack{i=1..n \\ rank(-Vaf_i)\le M}} I(q(i)>c)}\right)$$

**Zygosity of HLA alleles**

When an HLA allele is homozygous, we counted the neoantigens presented by that HLA allele by only once, not twice. The zygosity of HLA alleles will indeed affect the calculation of neoantigen load. However, it will be a lesser concern for CSiN. The calculation of CSiN is done in such a manner that it weighs whether truncal mutations generate more neoantigens or subclonal mutations generate more neoantigens. When an HLA allele is homozygous instead of heterozygous, the trend should be that it will affect the per-mutation neoantigen count of all mutations across the board. Therefore, this effect will tend to be cancelled out. However, there is another factor that might play into the effect of neoantigen repertoire found in each patient. When the two alleles of one HLA loci are the same, the same HLA proteins that bind the same neoantigen candidates will be translated. Depending on whether there are enough translated epitopes, the double HLA proteins may not have enough candidate epitope pool to bind. But when the HLA loci is heterozygous, the two alleles will likely bind differential epitope repertoire, thus avoiding this saturation effect. Therefore, it is hard to determine whether it is absolutely correct to count such neoantigens by once or by twice. Our current implementation only counts them once, though the user is welcome to finetune our R script for other possibilities.

**Ploidy and copy number variation**

The overall ploidy is a factor that influences all mutations and neoantigens, and thus is "cancelled out" in the calculation of CSiN for all mutations/neoantigens involved. CSiN is focused on determining the internal distributions of neoantigens to investigate whether more immunogenic neoantigens are concentrated in major tumor clones.

The calculation of VAF (#variant read/#total read) is influenced by copy number variation (CNV). If some of the tumor clones that have a particular mutation have, for example, copy number gain, then the VAF of this mutation in this tumor sample should be higher than when there is not any CNV. Higher CNV of a mutation will contribute to a higher expression level of the neoantigens translated from the gene hosting this mutation to some extent. And one can reasonably assume that the higher this expression level is, the more likely the neoantigen will have a stronger effect on the tumor cells with this mutation. Therefore, CNVs affect VAFs in the "correct" direction in terms of calculation of CSiN. But we welcome researchers to develop more advanced versions of CSiN that could possibly model CNV and VAF in a more sophisticated way.

**CSiN plot:**
We developed a specialized plot for intuitive visualization of the neoantigen clonal structure and how CSiN is calculated in each sample.

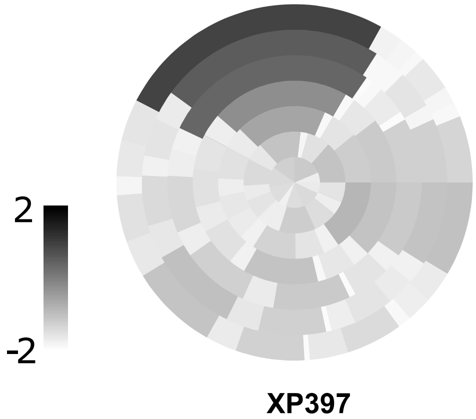


*Fig. S8 The CSiN plot for the primary tumor of XP397 from the UTSW KCP cohort is shown. The concentric circles from outermost to innermost are showing neoantigens satisfying increasingly stringent cutoffs on the strengths of binding, as are used in the definition of CSiN, to the MHC proteins. Mutations are shown in different "pies" of the circles, with area of one pie corresponding to the per-mutation neoantigen load. VAFs of the mutations are reflected as the coloring density of each "pie".*

**Validity of the neoantigen predictions**

We called the neoantigens of 6 melanoma patients from Ott *el al* (Ott et al. 2017). In this paper, the authors used genomics data to predict neoantigens using their pipeline, and picked 177 MHC class I neoantigens for experimental validation. 18 neoantigens were shown to be immunogenic by ELISPOT. We accessed their raw data, and used our pipeline to predict neoantigens. We examined, out of the neoantigens picked for experimental validation, how many can be found by our pipeline, and how many cannot be. We evaluated, out of these two groups of neoantigens, what proportion of neoantigens are immunogenic by EILSPOT standards. This was to test the specificity of our neoantigen prediction pipeline. We varied the RPKM threshold for neoantigen calling to test a variety of sensitivity levels. The results are shown in the following figure, which suggests that our neoantigen pipeline is slightly more specific (the called neoantigens are more likely to be positive by ELISPOT standards) than the neoantigen pipeline employed in the original study, given the same sensitivity level.

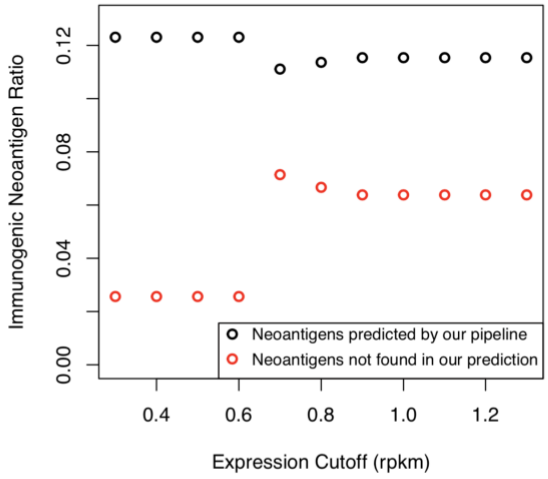


*Fig. S9 Validity of neoantigen predictions. The black dots stand for the portions of immunogenic neoantigens identified by our pipeline by ELISPOT standard. The red dots stand for the portions of immunogenic neoantigens not identified by our pipeline.*

Note: Only 18 out of all 177 neoantigens were shown to be immunogenic in the original study by ELISPOT standards. Regarding this issue, Ito *el al* (DOI:10.4172/2155-9899.1000322) examined multiple studies, and found that when researchers used the most common neoantigen validation experiment, ELISPOT, to validate neoantigen predictions from genomics data, the validation rate could go down to as low as 1%, in many cases. But this is very likely an underestimation due to many factors, such as that availability of matching TCRs happen to be extremely rare in the patients' sampled T cell repertoire for the neoantigen under examination.

**Driver of per-mutation neoantigen load**

For each individual mutation, the combination of mutation type, HLA alleles available in each patient, candidate neoantigens of all lengths, and candidate neoantigen sequences around the mutated positions (registers) will generate a pool of mutation-specific neoantigens. For each one of all the mutations in the same patient, the same HLA alleles and the same lengths (then naturally coupled with the same registers given the same length) will be "tried" to generate the whole pool of neoantigens for this mutation. So in this sense, they influence the per-mutation neoantigen load individually, but on the population average level, there is no difference for HLA allele, length, and register for mutations of high or low per-mutation neoantigen load.

The main driver of the number of neoantigens generated per mutation is the mutation type. The data presented below are from all patients analyzed in this study. It can be seen that frameshift mutations are likely going to generate the most neoantigens per mutation, while stoploss mutations also generate more neoantigens. Missense mutations and nonframeshift substitutions generate the lowest numbers of neoantigens per mutation. This observation is expected as insertions/deletions and stoploss mutations lead to the translation of completely new segments of protein sequences, compared to missense mutations and nonframeshift substitutions, which will generate neoantigens only in a short sliding window around the mutated position.

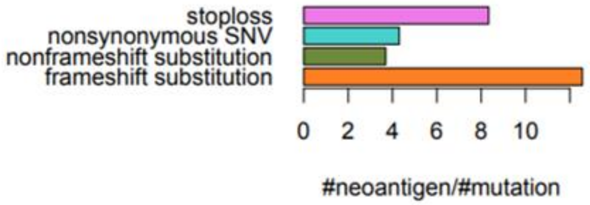


*Fig. S10 The average number of neoantigens generated by each type of mutations*

**<u>CSiN is independent of mutation load, neoantigen load, and transcriptomic-based biomarkers</u>**

We have shown the Spearman correlation between CSiN, mutation load, neoantigen load, and expression-based biomarkers in **Fig. 1D**. We also employed Pearson correlation, threshold comparisons, and mutual information to demonstrate the independence/dependence between these variables. In the following figure, Pearson correlation is used for (a), mutual information is used for (b). We set threshold as median of each variable in (c). Overall, our results suggest that CSiN is independent of these other variables.

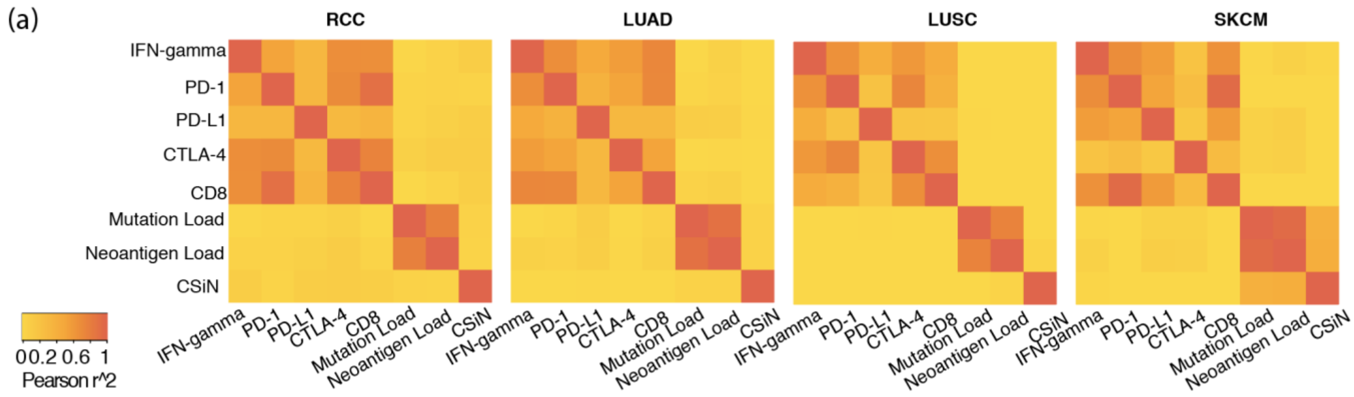

(a)
RCC
LUAD
LUSC
SKCM
IFN-gamma
PD-1
PD-L1
CTLA-4
CD8
Mutation Load
Neoantigen Load
CSiN
0 0.2 0.6 1
Pearson r^2


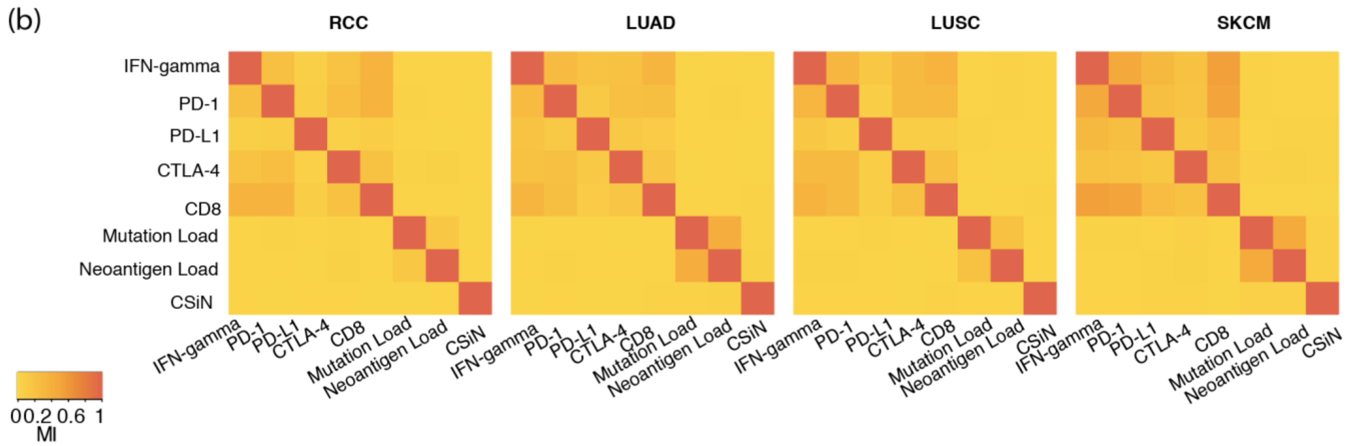

(b)
RCC
LUAD
LUSC
SKCM
IFN-gamma
PD-1
PD-L1
CTLA-4
CD8
Mutation Load
Neoantigen Load
CSiN
0 0.2 0.6 1
MI

(c)

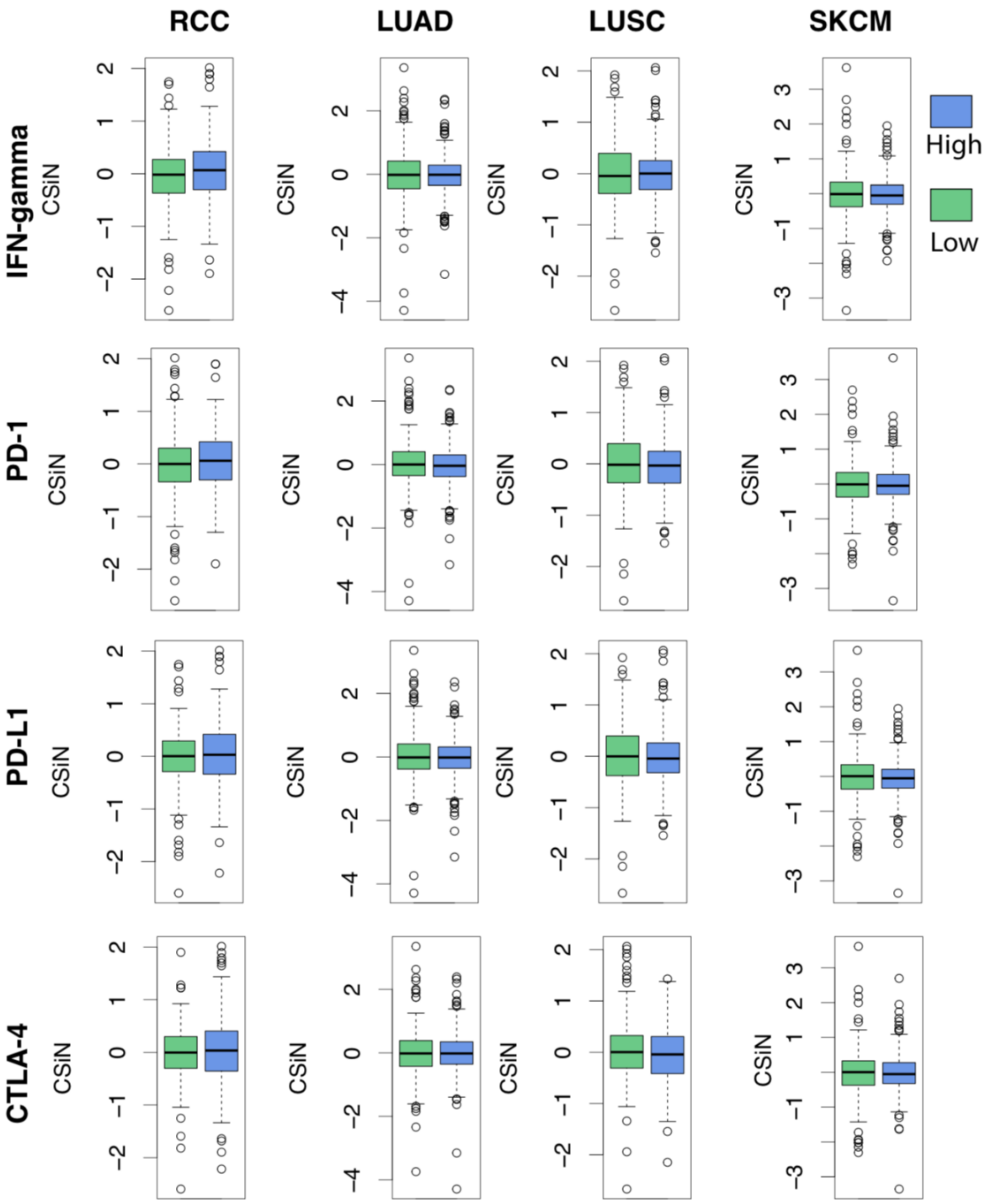

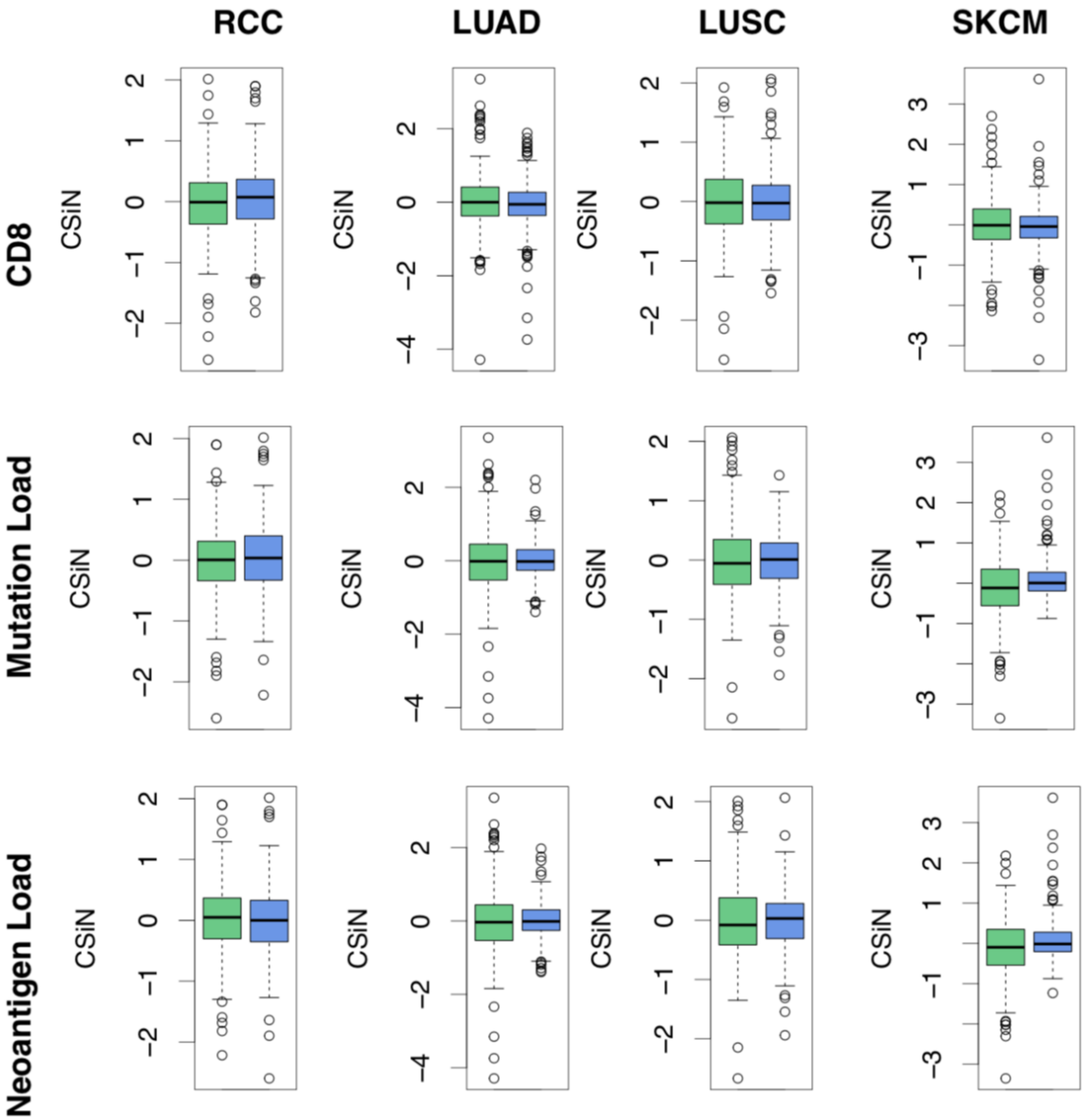


*Fig. S11 Demonstrating independence of CSiN from mutation load/neoantigen and transcriptomic-based biomarkers. (a) Pearson r square is used for pairwise correlation. (b) Mutual information is calculated for each two variables. (C) Median of each of the variable is set as threshold for CSiN threshold comparison.*

**<u>Comparing CSiN scores between primary and metastatic tumors</u>**

In our baseline survival cohorts (the patients shown in **Fig. 3**), we have annotations of which samples are primary tumors and which are metastatic samples. In the following figure (a), we showed that distant metastatic samples have a trend of decreasing CSiN compared with primary samples. These samples are not matched samples from the same patients. So we further identified a total of 7 patients from these cohorts that have genomics data available for their matched primary and metastatic samples. In (b), we showed that there is also a decreasing trend of CSiN in metastatic samples compared with primary samples. However, the P values for these comparisons are not significant and our conclusions are thus not definitive.

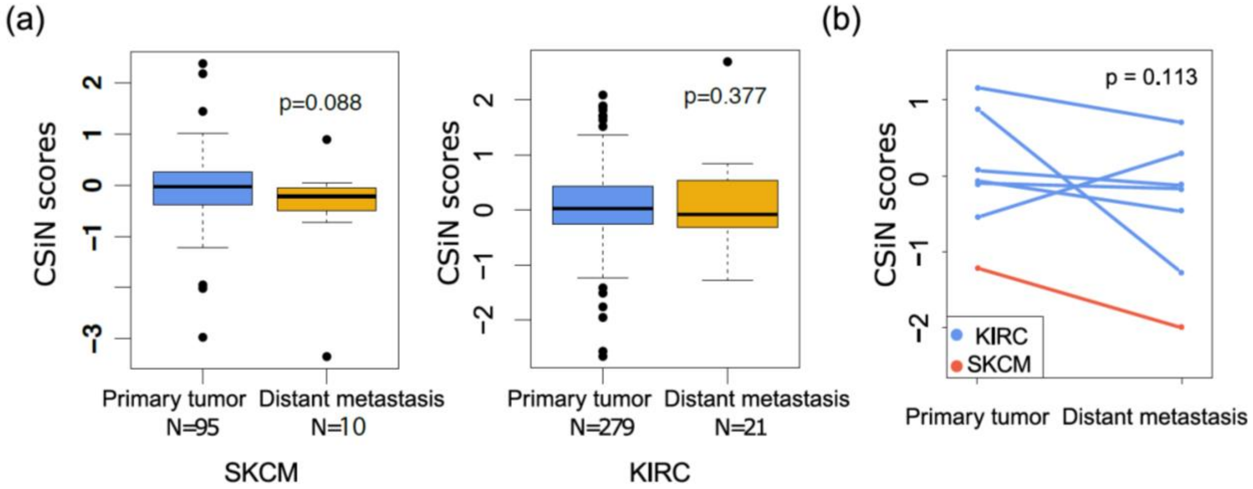


*Fig. S12 Association of CSiN with metastasis. (a) CSiN scores of RCC and SKCM patients without distant metastasis (N=279 and 95) and with distant metastasis (N=21 and 10) at time of biopsy. The first group includes primary tumors only. The second group includes samples from the primary sites or the distant metastasis sites, but all patients already had distant metastasis to another organ. (b) CSiN scores of 6 ccRCC patients and 1 melanoma patient with both matched primary tumor and distant metastasis genomics data available.*

**Evaluate the predictive power of CSiN to checkpoint inhibitor treatment using OS and PFS**

Overall survival (OS) data are available for the Riaz, Snyder, VanAllen, Hugo, and Miao cohorts. Progression-free survival (PFS) data are available for the Hellman and Rizvi cohorts. Here we show the predictive performance of CSiN, neoantigen load, and neoantigen fitness model, using OS/PFS as the criterion. Meta analyses of the Snyder, VanAllen, Hugo, Miao, and Hellman cohorts, through Fisher's method, yielded a fisher method for meta-analysis p value of 0.000563 for CSiN, 0.0706 for neoantigen load, and 0.0101 for the neoantigen fitness model.

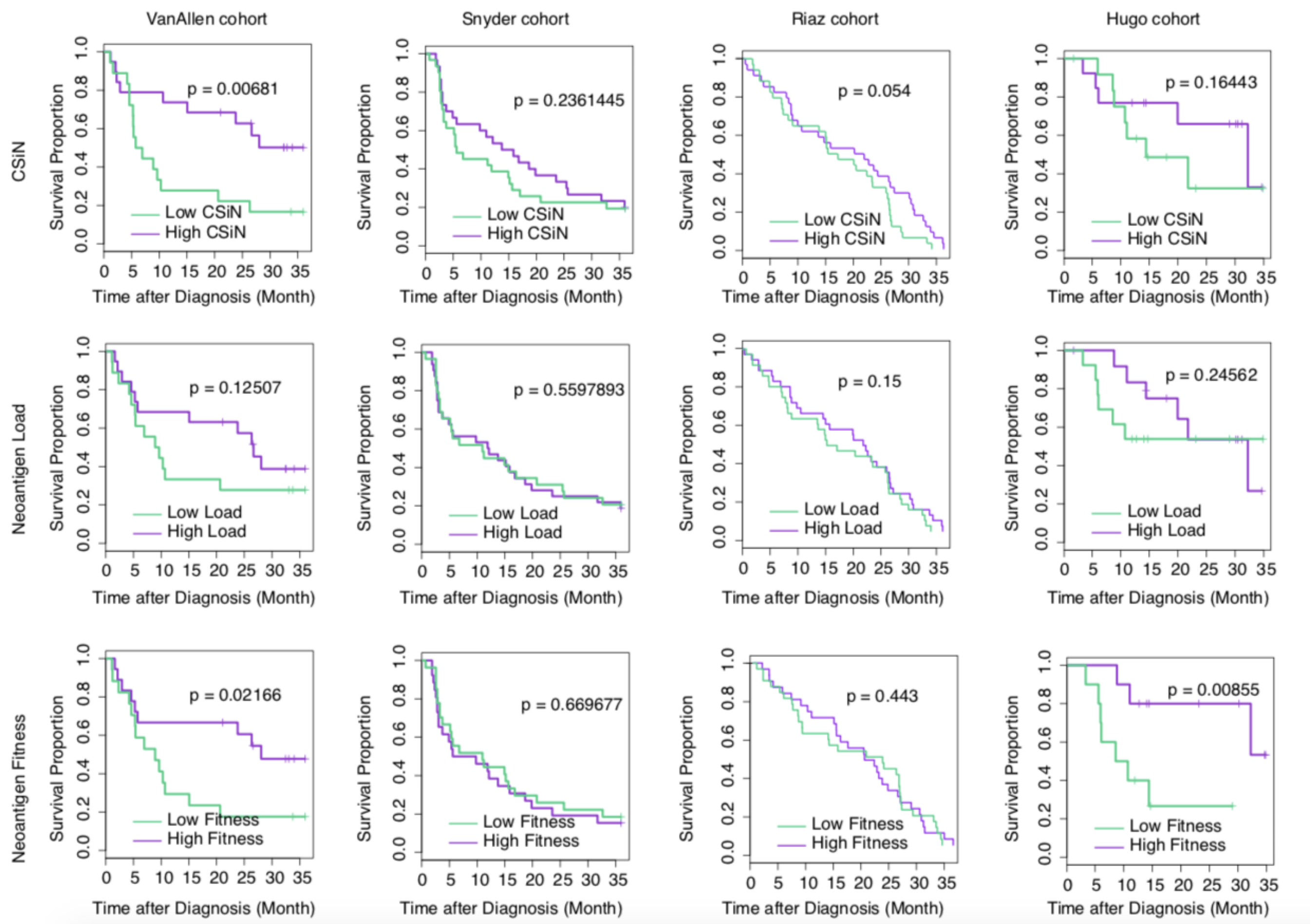

VanAllen cohort
Snyder cohort
Riaz cohort
Hugo cohort
CSiN
Neoantigen Load
Neoantigen Fitness
Survival Proportion
Time after Diagnosis (Month)
p = 0.00681
p = 0.2361445
p = 0.054
p = 0.16443
p = 0.12507
p = 0.5597893
p = 0.15
p = 0.24562
p = 0.02166
p = 0.669677
p = 0.443
p = 0.00855
Low CSiN
High CSiN
Low Load
High Load
Low Fitness
High Fitness

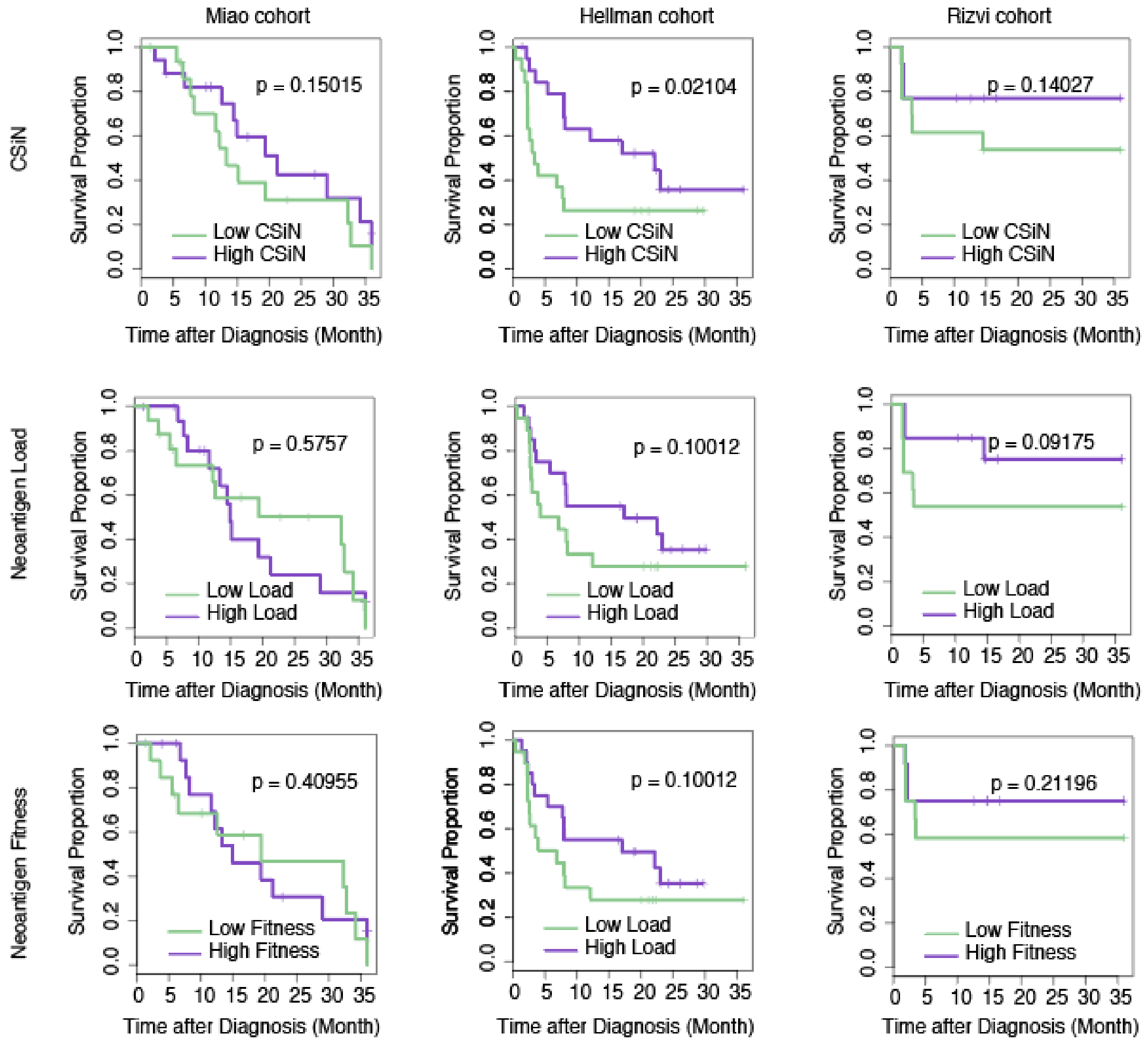


*Fig. S13 Validating the predictive power of CSiN, neoantigen load and neoantigen fitness model using OS/PFS as the criterion. Overall survival (OS) data are used for the Riaz, VanAllen, Hugo, and Miao cohorts. Progression-free survival (PFS) data are used for the Hellman and Rizvi cohorts.*

**Intra-tumor heterogeneity of CSiN and neoantigen loads**

On the UTSW KCP platform, we have done many multi-region samplings from the same individuals for a total of 39 patients and 121 samples (2 to 6 samples per patient). We analyzed those multi-region data, and show a comparison of the stability of CSiN and neoantigen load here (**Fig. 3**). We did analysis of variance for CSiN and neoantigen load for multi-region samples. F statistics, “between group variance (BGV)” over “within group variance (WGV)”, are comparable between CSiN and neoantigen load. Moreover, P values show that WGV is significantly smaller than BGV, for both neoantigen load and CSiN. Nevertheless, there is still some level of intra-tumor heterogeneity that can be observed in the multi-region sampling data of some patients, which demonstrates the challenges associated with using genomics-based biomarkers for clinical applications.

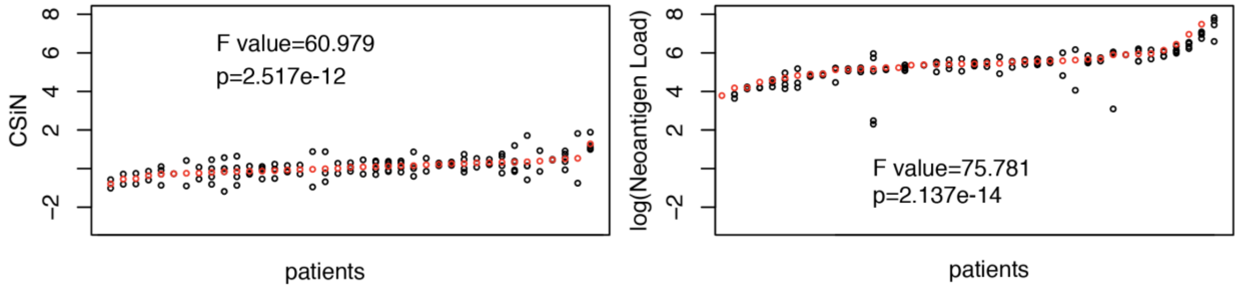


*Fig. S14 Assessing the intra-tumor heterogeneity of CSiN and neoantigen loads. Each black dot represents one sample. Dots with the same x coordinates stand for samples from the sample patient. Red dots stands for average values of the multi-region samples from the same patient.*

**Calculating CSiN with only exome-seq data**

The CSiN score can be calculated with exome-seq data only as a minimum. But we strongly prefer using RNA-seq data, if available. This will make the calculated CSiN score more accurate, as RNA-seq data can help filter out mutations in lowly expressed genes. Below, we show the results for all cohorts of **Fig. 2**, but we only used exome-seq data to calculate CSiN. As we had expected, the results are not as good as when we used RNA-seq data (when available) to filter the neoantigen lists for calculating CSiN. But importantly, most cohorts still remain statistically significant, and all cohorts uniformly show the same trend of better response correlated with higher CSiN scores.

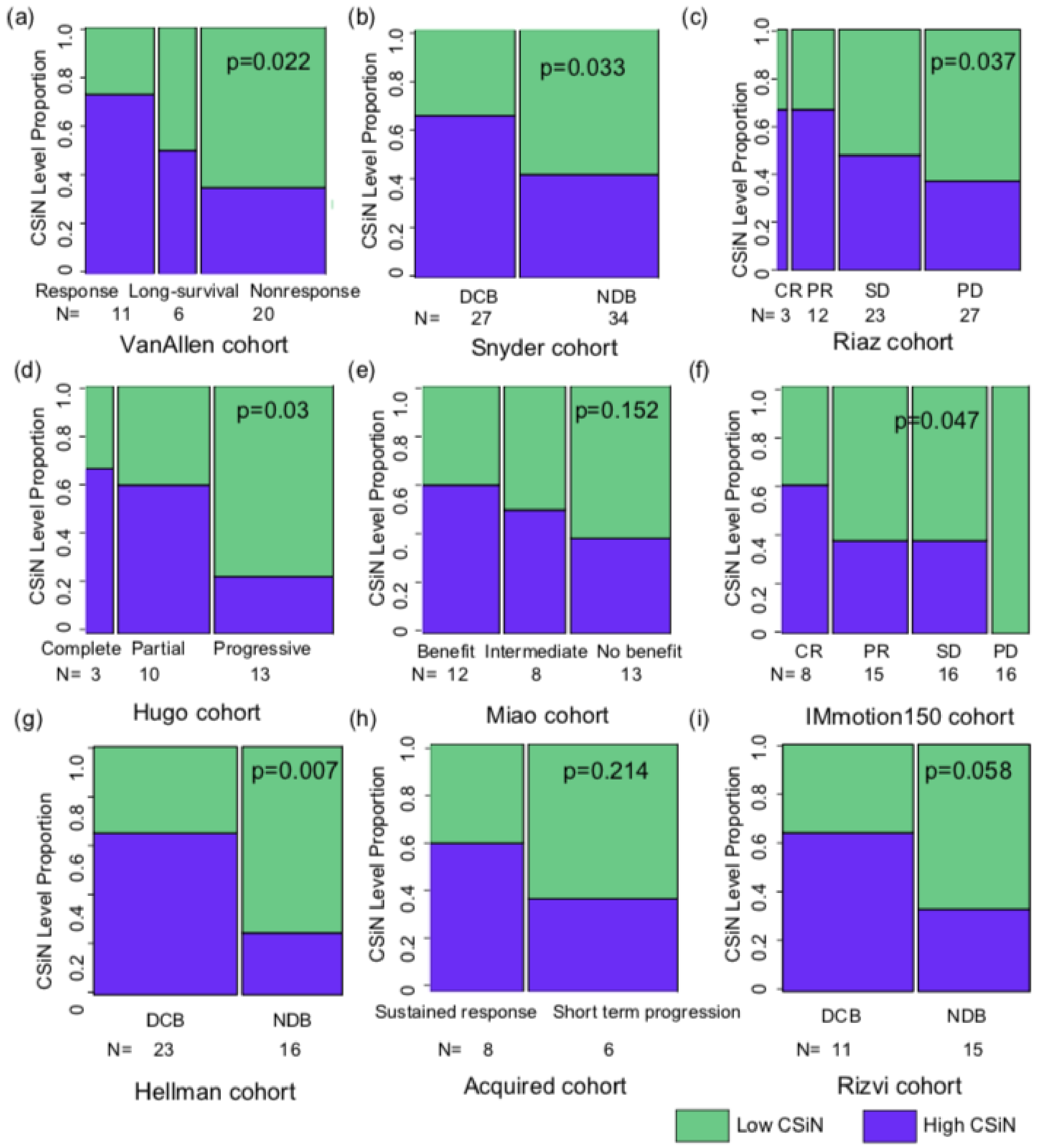


*Fig. S15 Calculating CSiN with only exome-seq data*

**<u>Using a more stringent cutoff on neoantigen load</u>**

For evaluating the predictive value of neoantigen load for immunotherapy treatment response, we also adopted another cutoff (median + 2 x interquartile range) that is more stringent than the median cutoff used in the main analyses, developed by Zehir *et al* (Zehir et al. 2017). We show the results here. However, the median + 2 x IQR cutoff has split the patients into two very unbalanced groups, and the high neoantigen load group has much fewer patients than the high neoantigen load group of patients based on median split, which may introduce instability due to small sample size.

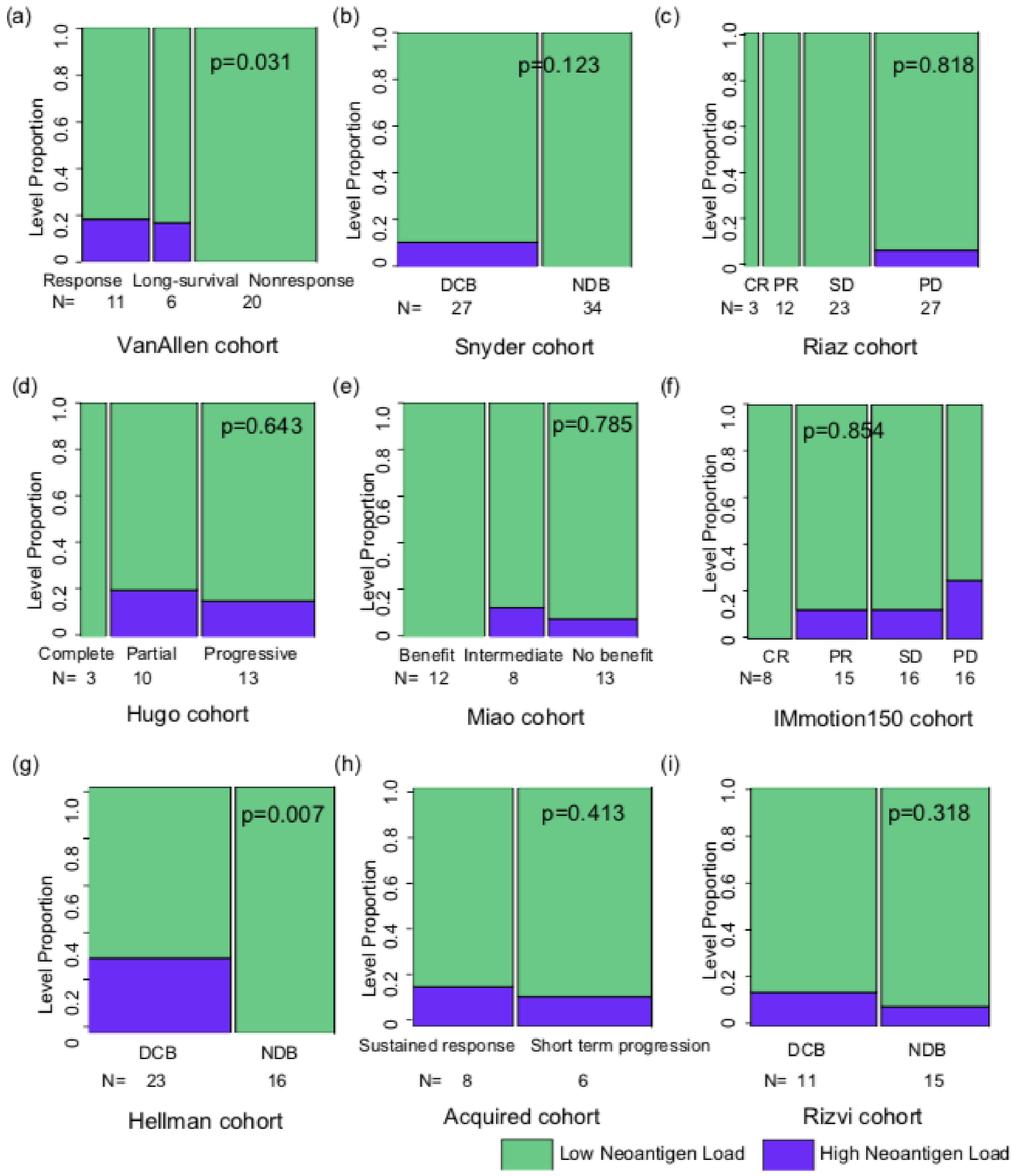


*Fig. S16 Using the median + 2 x interquartile range cutoff on neoantigen load*

**Separated analyses for class I and class II neoantigens**

For the 6 cohorts that were analyzed by our in-house pipelines, we have neoantigens of both class I and class II. For the other three cohorts, we don't have access to the raw genomics data, and we had to use the neoantigens called by the authors of the original reports. They happened to have only called class I neoantigens. For the first 6 cohorts, we calculated CSiN for class I and class II neoantigens, and showed the predictive power of the class I CSiN and class II CSiN separately. These class-specific CSiNs have less predictive powers for immunotherapy response (although for many cohorts, the trend of association is still the same and even statistical significance is attained in some cases), and this demonstrated the need for considering both class I and class II neoantigens in the calculation of CSiN.

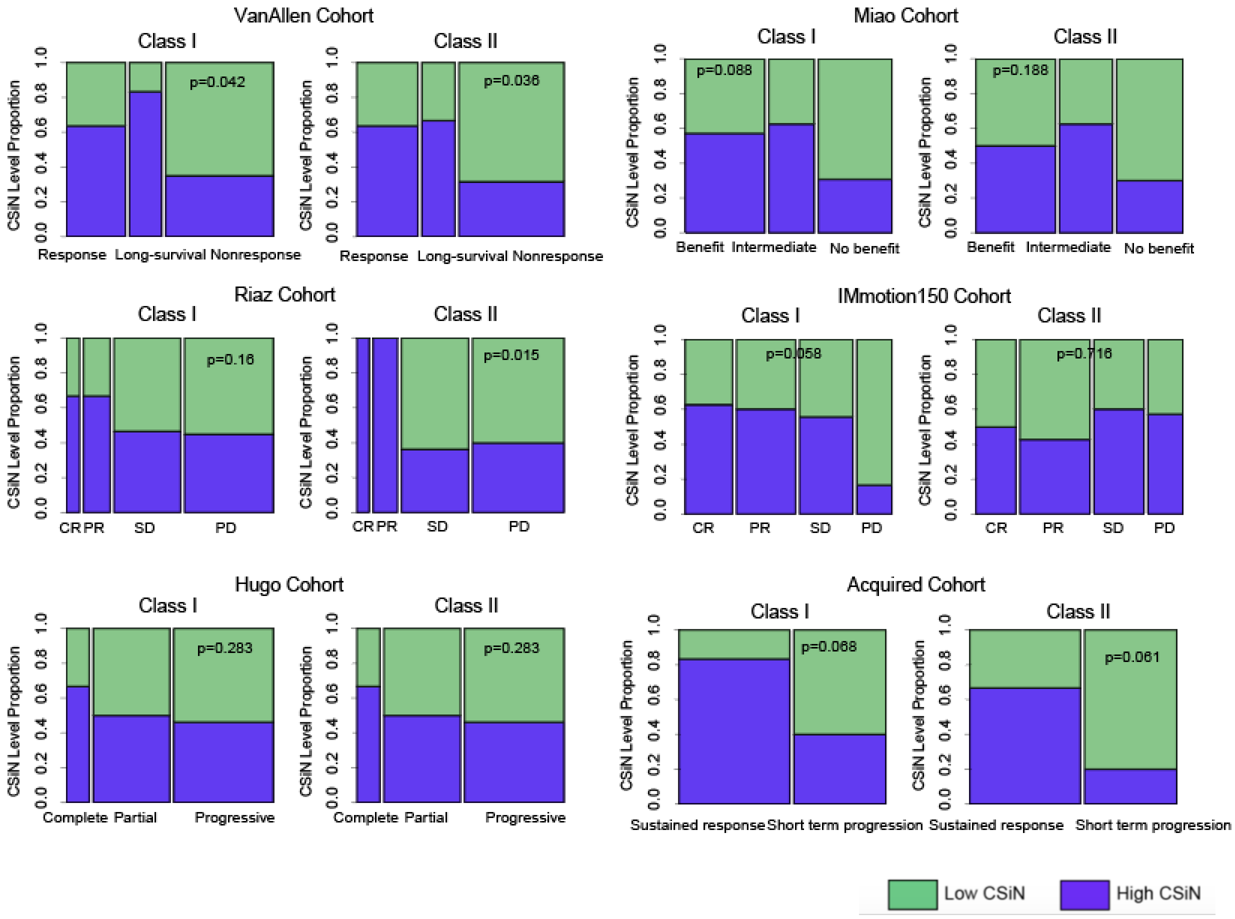


*Fig. S17 Predictive value of class I-specific CSiN and class II-specific CSiN*

For the neoantigen fitness model, we kept neoantigens that are missense, 9-mer, and class I as originally described in the neoantigen fitness study. We showed the association between neoantigen fitness and survival rate for the 3 cohorts that were also analyzed in the neoantigen fitness paper. The result is largely consistent with the original neoantigen fitness study, with slightly larger p values, probably due to a number of differences in data pre-processing (mutation calling, neoantigen calling, *etc*) that exist between their study and own study.

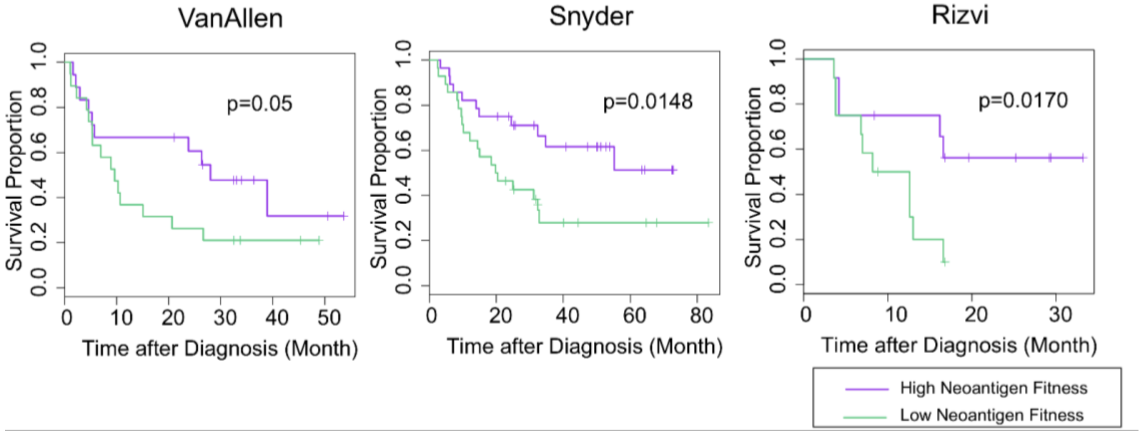


*Fig. S18 Predictive value of class I-specific neoantigen fitness model measured by survival analyses (limiting to 9-mers from missense mutations)*

We also kept missense, 9-mer, and class I neoantigens for calculating the neoantigen fitness model for the other 6 cohorts, and presented the predictive power of neoantigen fitness using the categorical response variable used in the main analyses, for all 9 cohorts. The results are shown below. We observed that the neoantigen fitness showed good association with patients' responses in three out of all 9 cohorts.

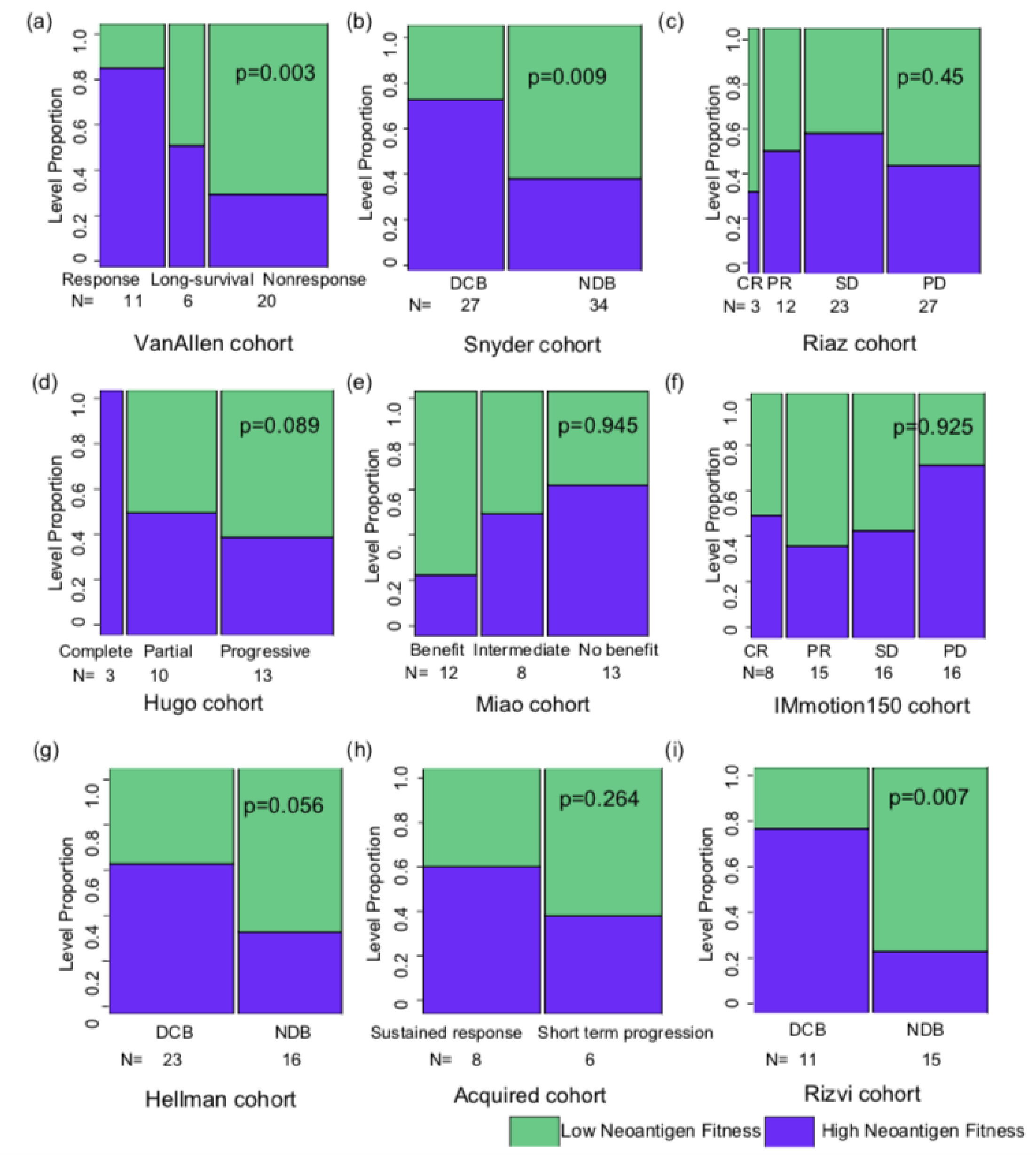


*Fig. S19 Predictive value of class I-specific neoantigen fitness model measured by categorical response variables (limiting to 9-mers from missense mutations)*

### Testing correlation between treatment response and CSiN in the $T_{eff}$-high subset

In **Fig. 2F**, we performed stratified analyses and showed that CSiN is predictive of treatment response in the $T_{eff}$-high subset of the IMmotion150 cohort. Here we also subset the VanAllen, Riaz, Hugo and Miao cohorts that have RNA-Seq data available for calculating $T_{eff}$ signature expression, and showed the predictive value of CSiN in the $T_{eff}$-high (60%) subsets.

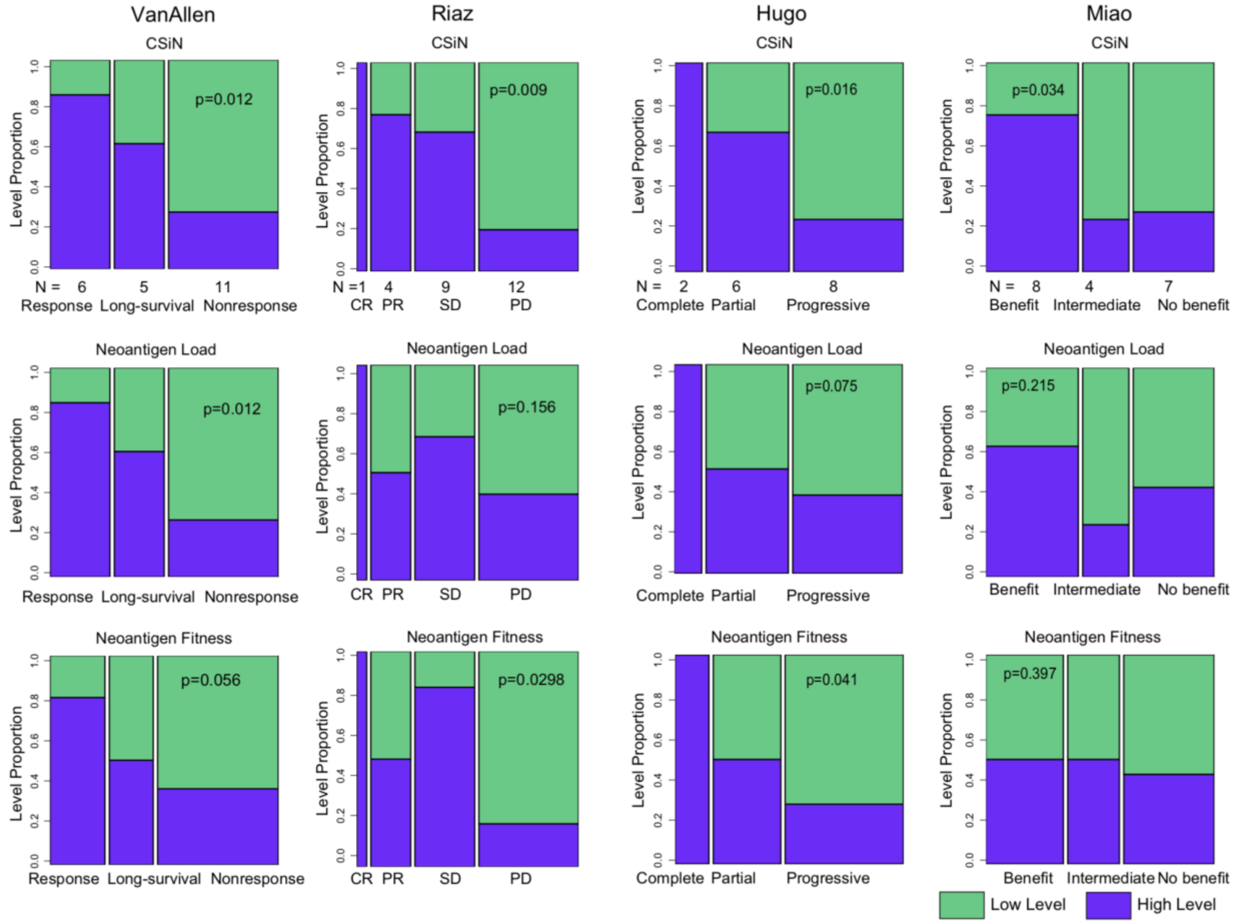


*Fig. S20 Predictive value of CSiN for the patients with high $T_{eff}$ signature expression. For the Riaz cohort, only a subset of the patients have matched RNA-Seq data. So the 60% for this cohort was selected from these patients only.*

**Testing correlation between treatment response and CSiN for all the patients in the IMmotion150 cohort**

In the following figure, we showed the correlation between treatment response and CSiN for all the patients who received Atezolizumab and all the patients who received Sunitinib, in the IMmotion150 cohort. No subsetting based on $T_{eff}$ expression was carried out as in Fig. 2

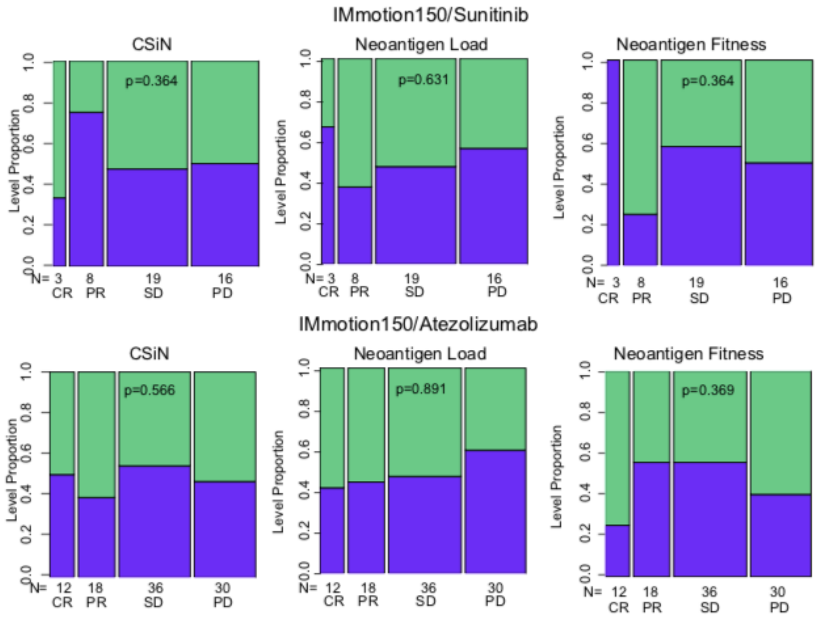


*Fig. S21 Predictive value of CSiN for the patients treated by sunitinib and by atezolizumab in the IMmotion150 cohort.* $T_{eff}$ *high and low patients are all included.*

**Testing correlation between treatment response and CSiN for all the patients in the Hellmann cohort**

In the following figure, we showed the correlation between treatment response and CSiN for all the patients in the Hellmann cohort. No subsetting based on PD-L1 expression was carried out as in **Fig. 2**

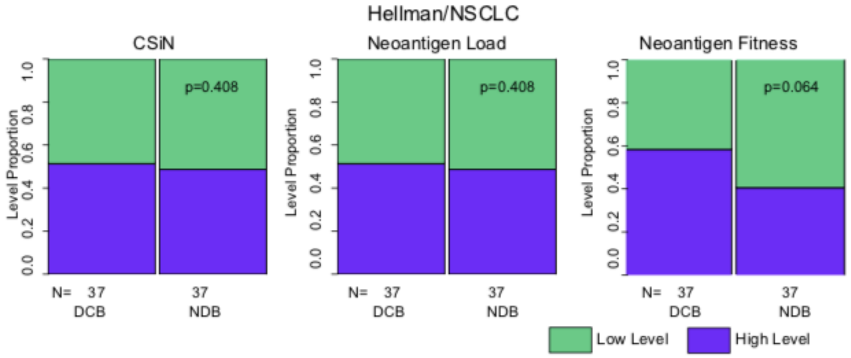


*Fig. S22 Predictive value of CSiN for all the patients in the Hellmann cohort*

**CSiN vs. tumor heterogeneity**

We didn't observe a correlation between CSiN and tumor heterogeneity. Here we show that the number of tumor clones determined by PyClone and CSiN score is not correlated. The pearson correlation for each type of cancer is 0.034 (KIRC), 0.018 (LUAD), -0.00052 (LUSC), and 0.112 (SKCM). This was also shown in boxplots below.

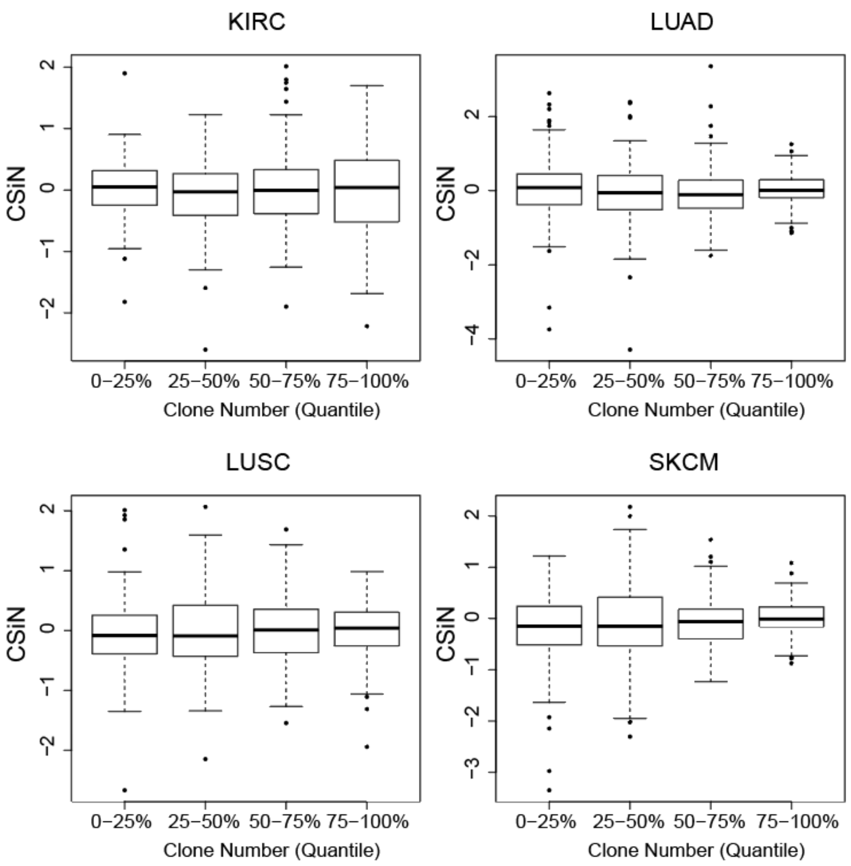


*Fig. S23 Boxplots showing distribution of CSiN scores in quartiles of tumor clone number determined by pyclone*
